# Phylogenomic patterns of divergence and gene flow detail the evolution of reinforcement and hybrid speciation in *Phlox* wildflowers

**DOI:** 10.1101/2022.04.15.488502

**Authors:** Austin G. Garner, Benjamin E. Goulet-Scott, Robin Hopkins

**Author notes:** Authors to whom correspondence should be addressed: Austin G. Garner, Robin Hopkins.

## Abstract

- The tree of life is riddled with reticulate evolutionary histories, and some clades, such as the eastern standing *Phlox*, appear to be hotspots of hybridization. In this group there are two cases of reinforcement and nine hypothesized hybrid species. Given the historical importance of this group for our understanding of plant speciation, the relationships of these taxa and the role of hybridization and gene flow in their diversification require genomic validation.
- Using phylogenomic analyses, we resolve the evolutionary relationships of the eastern standing *Phlox* and test hypotheses about if and how hybridization and gene flow were creative forces in their diversification.
- Our results provide novel resolution of the phylogenetic relationships in this group, including well-supported paraphyly across some species. We identify gene flow during one of two cases of reinforcement and find support for one of the five hypothesized homoploid hybrid speciation events. Additionally, we infer the ancestries of four allotetraploid hybrid species.
- Hybridization has contributed to diverse evolutionary outcomes within this *Phlox* group; although, not as extensively as previously hypothesized. This study demonstrates the importance of phylogenomics in confirming hypothesized histories of non-model systems and adds to the growing evidence of interspecific genetic exchange in the generation of biodiversity.

## Introduction

Hybridization and gene flow can play important and diverse roles during the formation and divergence of species (Taylor & Larson, 2019). While these processes are often viewed as homogenizing forces, they can also generate novel variation. For example, gene flow can create stable hybrid species and/or fuel the diversification of adaptive radiations (Abbott *et al*., 2013; Schumer *et al*., 2014; Marques *et al*., 2019). The act of hybridization itself can also create selective pressures favoring the evolution of reproductive trait divergence to reduce mating between species, i.e. reinforcement (Howard, 1993; Garner *et al*., 2018). The effects of hybridization and gene flow on speciation have been investigated across the tree of life, yet we lack a broader understanding of the frequency and extent to which these forces impact speciation across clades of organisms (but see Eaton *et al*., 2015; Pease *et al*., 2016). Characterizing the context and consequences of hybridization and gene flow across groups of species is necessary for discerning how these forces influence the generation and maintenance of biodiversity.

Much of the foundational work on hybridization as a creative force in speciation comes from historic research on non-model plant systems (Anderson, 1949; Stebbins, 1959; Grant, 1981). Patterns of morphological, physiological, and ecological trait variation across and within plant species created taxonomic conflict, inspiring biologists to propose some species arose from interspecific mating (Goulet *et al*., 2017). In particular, observations of populations with combinations of intermediate, recombinant, and novel traits to those of nearby geographically overlapping species inspired hypotheses of polyploid and homoploid hybrid speciation (Gottlieb, 1972; Soltis & Soltis, 2009; Yakimowski & Rieseberg, 2014; Barker *et al*., 2016). It is important to return to these known and hypothesized cases of reticulate evolution with genomic tests for gene flow to confirm these evolutionary processes and further understand the context of their evolution (Goulet *et al*., 2017).

Similarly, patterns of trait divergence in sympatry, where species coexist, but not in allopatry, motivated hypotheses of hybridization causing reproductive trait divergence by reinforcement (Hopkins, 2013). Yet, only a few empirical studies have evaluated the presence and extent of gene flow during the evolution of reinforcement in nature (Kulathinal *et al*., 2009; Roda *et al*., 2017; Turissini & Matute, 2017; Lemmon & Juenger, 2017; Dyer *et al*., 2018). Maladaptive hybridization can drive selection for reinforcement and the divergence of plant reproductive traits such as flower color, flowering time, and cross-compatibility (McNeilly & Antonovics, 1968; Levin, 1985; Fishman & Wyatt, 1999). However, gene flow and recombination can also impede divergence by reinforcement (Felsenstein, 1981). Much debate and theory has focused on if and how reinforcement can evolve if hybridization also causes interspecific gene flow (Servedio & Noor, 2003), but further genomic study of cases of reinforcement are needed to understand if and how this reproductive trait divergence has evolved with gene flow.

Hybridization and gene flow result in the mixing of genetic variation between species. If these processes coincide with the formation of novel diverging lineages, admixture of parental species’ genetic variation will be present in the new lineage. Traditional phylogenetic methods infer evolution along bifurcating trees. However, modern phylogenomic analyses can use discordance in allele patterns under these trees (Green *et al*., 2010; Durand *et al*., 2011) or model allele frequencies at the tips (Gutenkunst *et al*., 2009; Pickrell & Pritchard, 2012; Excoffier *et al*., 2013) to identify a history of gene flow between taxa (reviewed in Hibbins & Hahn, 2021). Contemporary genome-wide sequencing methods, such as restriction site-associated DNA (RAD) sequencing (Andrews *et al*., 2016), allow cost-efficient application of these methods to infer the evolutionary histories of genetic exchange in non-model systems (Eaton & Ree, 2013; Eaton *et al*., 2015; Léveillé-Bourret *et al*., 2020; Bombonato *et al*., 2020; Guo *et al*., 2021; Suissa *et al*., 2022).

The eastern standing *Phlox* (Polemoniaceae) wildflowers are an ideal system for exploring the role of hybridization and gene flow in speciation. They are a monophyletic group (Landis *et al*., 2018) distinct from the rest of *Phlox* by their upright growth habit and natural occurrence across mid to eastern North America. Their natural ranges overlap and intertwine, generating numerous zones of species contact (Fig. 1), and reproductive barriers between these taxa are highly permissive (Levin, 1966b). Decades of morphological, ecological, and biochemical evidence have suggested hybridization and gene flow between taxa, leading to hypotheses of their diversification by hybrid speciation and reinforcement (Anderson & Gage, 1952; Erbe & Turner, 1962; Levin, 1966a; Levin & Smith, 1966; Levin & Kerster, 1967; Levin & Schaal, 1970b; Levin, 1985).

**Figure 1.**
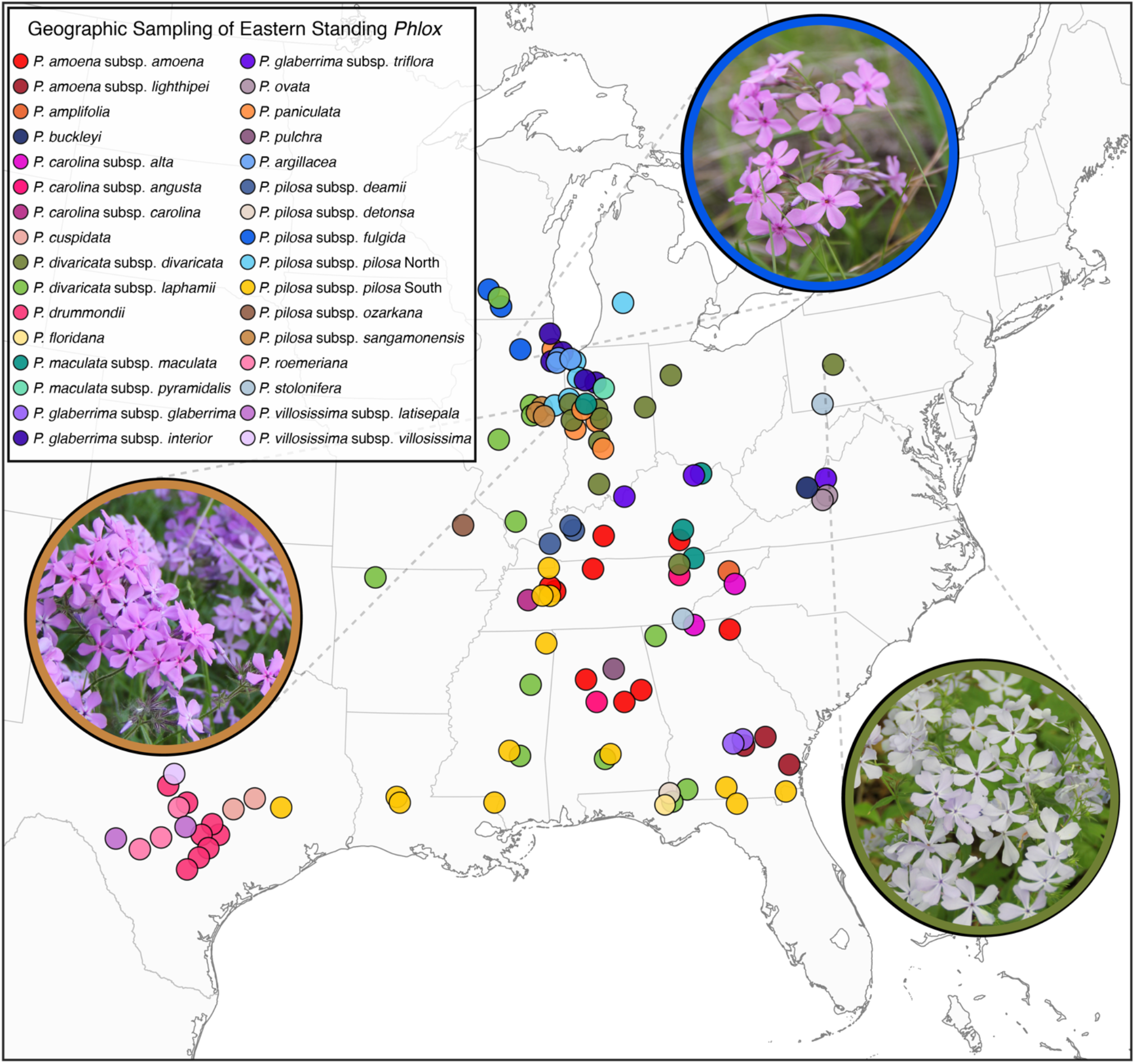
Map of individuals sampled across the interdigitating ranges of the thirty-two eastern standing *Phlox* taxa included for phylogenetic analyses. Each dot represents the locality of a single individual with the color corresponding to an individual’s taxonomic identity. Three insets show *P. pilosa* subsp. *sangamonensis* (brown), *P. pilosa* subsp*. fulgida* (blue), and *P. divaricata* subsp. *divaricata* (green) in their natural habitat.

There are five hypothesized cases of homoploid hybrid species (*P. argillacea, P. maculata* subsp. *pyramidalis, P. pilosa* subsp. *deamii, P. pilosa* subsp. *detonsa*, and *P. amoena* subsp. *lighthipei*) and at least four cases of allotetraploid hybrid species (*P. villosissima* subsp. *villosissima, P. villosissima* subsp. *latisepala, P. floridana, P. buckleyi*) (Table 1) within the eastern standing *Phlox*. Support for these hypotheses were originally based on observations that putative hybrid lineages had reproductive, morphological, and ecophysiological trait values that appear to be intermediate, recombined, or transgressive with putative parental taxa, making them potentially reproductively and ecologically distinct from their parental lineages (Levin, 1963, 1969; Levin & Smith, 1966; Hadley & Levin, 1969). Further analysis of karyology (Smith & Levin, 1967; Levin, 1968), cross compatibility (Levin, 1966b), biochemical profiles (Levin & Schaal, 1970b, 1972; Levy & Levin, 1974, 1975), and microsatellite markers (Fehlberg *et al*., 2014) built support for these hypotheses. Finally, natural hybrid zones and synthetic crosses between pairs of putative parental taxa generated hybrids with phenotypes similar to the putative stable hybrid lineages in nature (Levin, 1963, 1966a; Levin & Smith, 1966). However, other treatments of these putative hybrids species have assumed them to be diverging varieties without hybridization defining their formation (Wherry, 1956) or found no support for hybrid ancestry with genome-wide markers (Goulet-Scott *et al*., 2021).

**Table 1.**
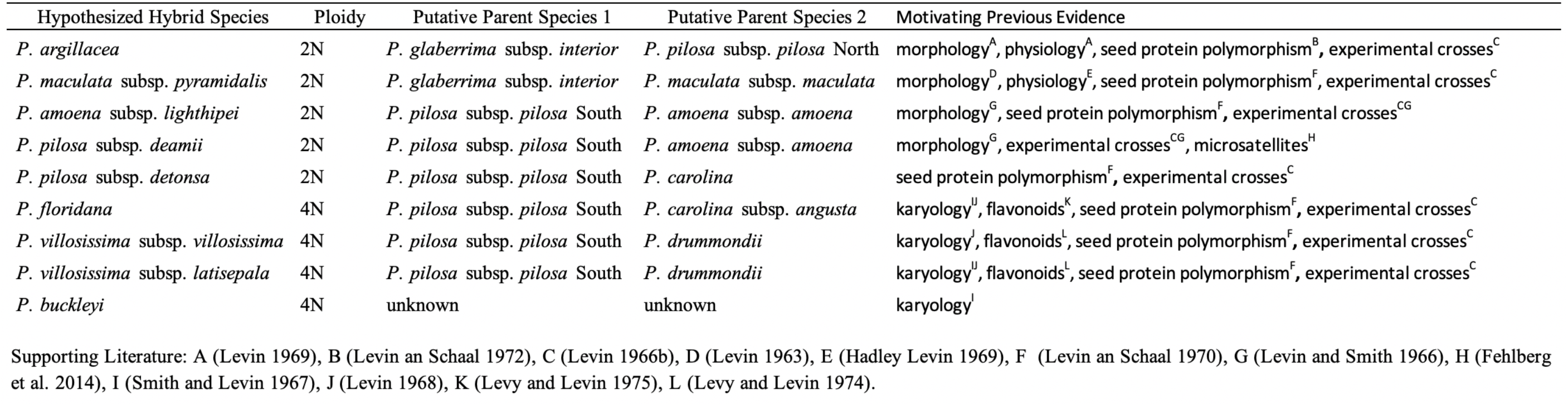
Hybrid speciation hypotheses in the eastern standing *Phlox*

This clade of *Phlox* also has two cases of reinforcement. During the speciation of both *P. drummondii* and *P. pilosa* subsp. *pilosa*, flower color divergence within sympatric populations decreases hybridization with closely related species (reviewed in Hopkins, 2013; Fig. 3). *P. drummondii* has evolved from having light-blue to dark-red flower color to prevent hybridization with sympatric light-blue colored *P. cuspidata* (Levin, 1985; Hopkins & Rausher, 2012), and *P. pilosa* subsp. *pilosa* have evolved from pink to white flower color to prevent hybridization where it cooccurs with pink colored *P. glaberrima* subsp. *interior* (Levin & Kerster, 1967). Biochemical evidence suggests gene flow occurred between sympatric *P. pilosa* subsp. *pilosa* and *P. glaberrima* subsp. *interior* (Levin & Schaal, 1972), and genomic analyses have inferred a history of gene flow between sympatric *P. drummondii* and *P. cuspidata*, but analyses were limited to single individuals (Roda *et al*., 2017).

**Figure 2.**
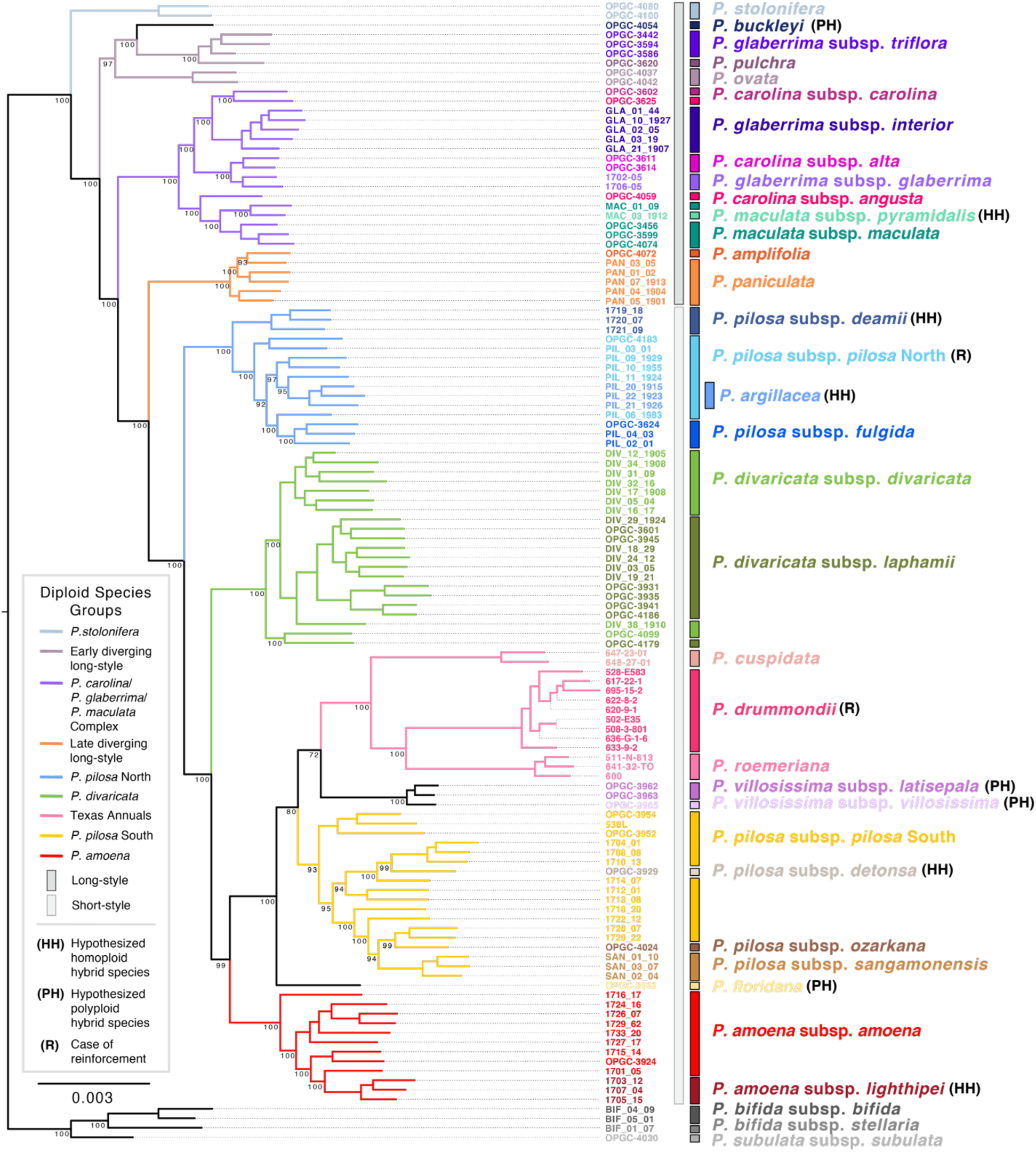
Maximum likelihood phylogenetic inference of eastern standing *Phlox* from whole concatenated ddRAD loci. Loci were required to be shared by at least 30 individuals. Numbers on branches represent bootstrap supports from 100 bootstrap replicates using ultrafast bootstrap approximation (UFB). Bootstrap supports are shown for all nodes differentiating between taxa.

**Figure 3.**
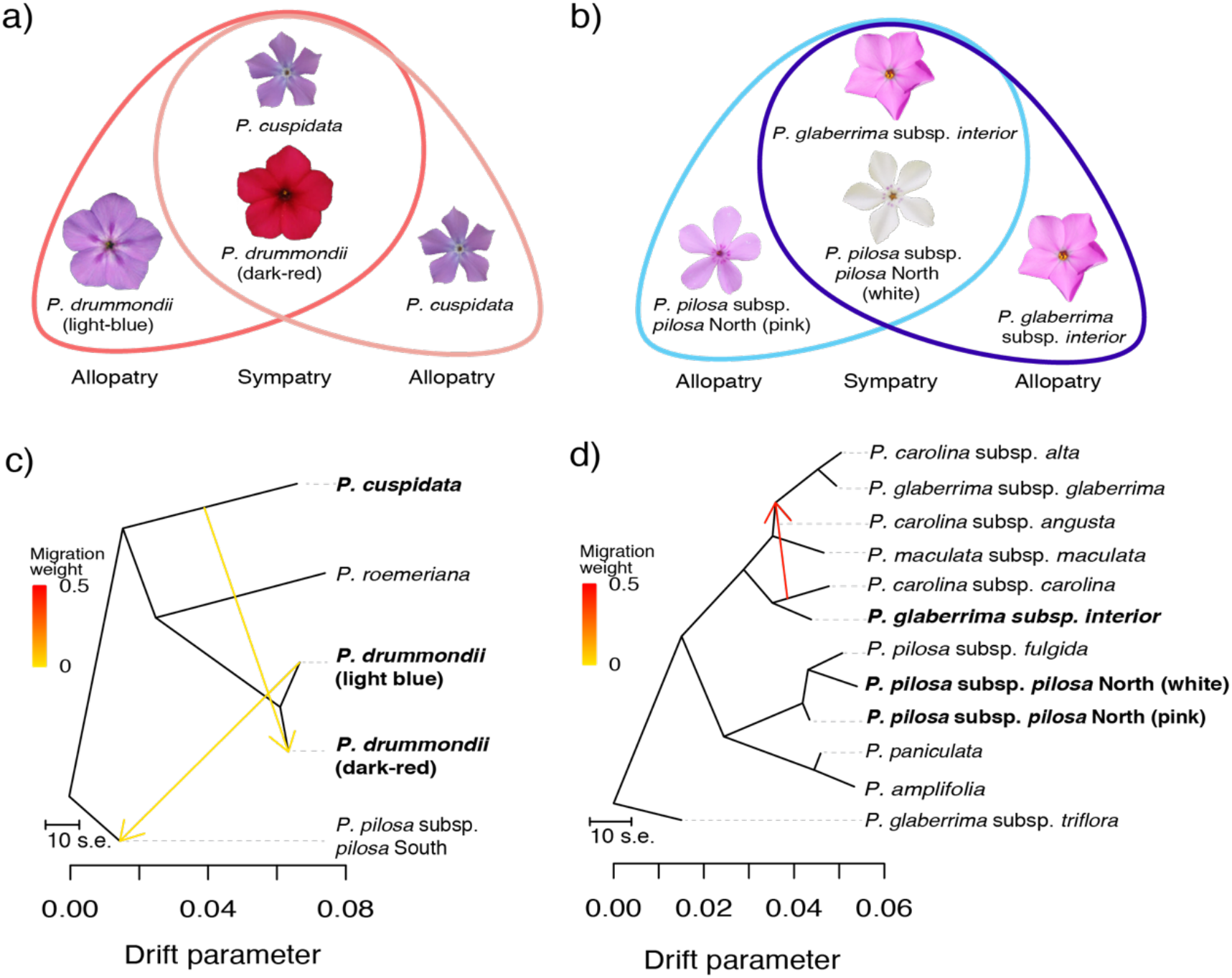
(a and b) Schematics of flower color divergence due to reinforcement in two species of *Phlox* to prevent maladaptive hybridization with another species in sympatry. (a) *P. drummondii* (red) has similar light-blue colored flowers to *P. cuspidata* (pink) in allopatry but *P. drummondii* has evolved dark-red flowers in sympatry to reduce hybridizing with *P. cuspidata*. (b) In allopatry *P. pilosa* subsp. *pilosa* North (light blue) has similar pink colored flowers to *P. glaberrima* subsp. *interior* (dark blue), but where they co-occur in high frequency in sympatry, *P. pilosa* subsp. *pilosa* North has evolved white flowers to reduce hybridizing with *P. glaberrima* subsp. *interior*. (c and d) Best fit models from TreeMix inferred gene flow between *P. cuspidata* and dark-red *P. drummondii* (c) (best fit m=2) but did not support evidence of gene flow between *P. glaberrima* subsp. *interior* and either color morph of *P. pilosa* subsp. *pilosa* North (d) (best fit m=1).

*Phlox* has been a taxonomically and phylogenetically difficult genus. Traditional delineations of *Phlox* species relied on morphological, ecological, and geographic characteristics, resulting in a long history of describing, promoting, and demoting species, subspecies, and varieties (Whitehouse, 1945; Wherry, 1955, 1965; Erbe & Turner, 1962; Turner, 1998; Locklear, 2011a,b). Conspicuous variation in pistil style length has been used to divide the eastern standing *Phlox* into long-styled (10-26mm) and short-styled (1.5-4mm) species groups (Wherry, 1930, 1931, 1932, 1955; Ferguson *et al*., 1999). However, other morphological traits vary widely within and between species, and species-level relationships within the group remain either poorly supported or incongruent across phylogenetic studies (Ferguson *et al*., 1999; Ferguson & Jansen, 2002; Landis *et al*., 2018; Goulet-Scott *et al*., 2021). Resolving the phylogenetic relationship of the eastern standing *Phlox* with modern genomic methods will provide a powerful framework for returning to evolutionary hypotheses about their speciation and understanding the dynamics by which their diversification has occurred.

Here, we apply double-digest RAD (ddRAD) sequencing with phylogenomic approaches to resolve the evolutionary relationships of the eastern standing *Phlox*. We leverage this phylogenetic framework to examine the contribution of hybridization and gene flow in the diversification of this system. Specifically, we test if gene flow has occurred alongside the evolution of two cases of reinforcement and investigate patterns of genetic composition and gene flow in multiple hypothesized cases of homoploid and allotetraploid hybrid speciation.

## Materials and Methods

### Sampling, sequencing, and data processing

We collected wild plants from across the native ranges of the eastern standing *Phlox* from 2017-2019, and we acquired additional samples from natural populations preserved in tissue culture at the Ohio State University Ornamental Plant Germplasm Center (OPGC). Our sampling includes seventeen recognized eastern standing *Phlox* species, representing thirty-two described subspecies, and three outgroup taxa from the mat-forming *Phlox* (Fig. 1; Supporting Information Table S1). We split taxa following Locklear, 2011b, except for the Texas annuals for which we only designated at species level.

Plants were grown in the greenhouse under controlled conditions until flowering. We extracted genomic DNA from bud tissue using a hybrid Omega Bio-Tek EZNA DNA kit and CTAB/Chloroform extraction protocol. Genomic libraries were constructed with a double-digest restriction-site-associated digestion (ddRAD) sequencing protocol and paired-end sequenced as described in Goulet-Scott *et al*., 2021. Raw sequence reads were demultiplexed, filtered, and *de novo* assembled as described in Goulet-Scott *et al*., 2021. Samples with minimal informative shared loci were removed, resulting in a final dataset of 116 ingroup and 4 outgroup individuals.

For the 120 retained samples, filtered raw reads (average of 3,060,421 reads per individual) were processed by iPyRAD v.0.9.50 (Eaton & Overcast, 2020) to construct four datasets with varying levels of missingness, requiring a locus to be shared among a minimum of N = 4, 10, 20, and 30 individuals, named MS4, MS10, MS20, and MS30 respectively. These assembled datasets produced matrices with a total of 165,549 to 5,3871 loci and 1,884,423 to 105,363 SNP sites with a locus missingness rate of 90%, 83%, 72%, and 60% respectively.

### Phylogenetic inferences

Phylogenetic relationships of our samples were inferred using concatenation and coalescent-based methods. First, whole ddRAD loci from the MS10, MS20 and MS30 datasets were individually concatenated into supermatrices. Maximum likelihood phylogenies were estimated from each supermatrix using IQ-TREE v.1.6.10 (Nguyen *et al*., 2015) with the GTR + gamma nucleotide substitution model and 1000 bootstrap replicates using ultrafast bootstrap approximation (Minh *et al*., 2013; Hoang *et al*., 2018).

Second, we inferred a population-level species tree using SVDquartets (Chifman & Kubatko, 2014), as implemented in PAUP v.4.0a166 (Swofford, 2002). This method uses unlinked SNPs to infer relationships among quartets of taxa under a coalescent model and then estimates a species tree using a quartet assembly method. We conducted the analysis on our dataset with least missingness, MS30, with the mat-forming species removed and *P. stolonifera* set as the outgroup. To avoid genetic linkage, we sampled a single SNP-site with the least amount of missing data from each locus, totaling 5,306 variant sites. We used all 7,160,245 possible quartets and 100 non-parametric bootstrap replicates to assess topological support and to generate a majority-rule consensus species tree.

Support values between these two inference methods are derived from different estimation methods and cannot be compared one to one. Standard bootstrap supports (SBS), as used with our SVDquartets inferred tree, underestimate the likelihood of relationships; SBS supports of 80% have a 95% likelihood of being correct. Ultrafast bootstrap supports (UBS), as used for our concatenation inference method, are unbiased when >70%, so a 95% support has a 0.95 probability of being correct (Minh *et al*., 2013).

### Tests for gene flow alongside reinforcement

We tested for evidence of gene flow alongside the evolution of reinforcement in *P. drummondii* and *P. pilosa* subsp. *pilosa*. First, we computed Patterson’s D-statistic (Green *et al*., 2010; Durand *et al*., 2011) to infer if gene flow occurred between two lineages in a given phylogeny. This test measures the asymmetry in the ratio of two discordant allele patterns, ABBA and BABA, across the topology of a four-taxon pectinate tree ((P1, P2), P3), O). Under stochastic lineage divergence without gene flow between lineages P3 and P1 or P3 and P2, the proportion of discordant allele patterns from incomplete lineage sorting are unbiased (ABBA=BABA); however, when introgression has occurred between P3 and P1 or P3 and P2, one discordant allele pattern is more prevalent than the other (ABBA>BABA, BABA>ABBA).

We defined the P1, P2, P3, and O lineages as taxa and a “test” as a unique combination of individuals from these taxa. We subset the complete MS4 data matrix to include the individuals from a set of four taxa that fit the pectinate topology under our ML phylogenetic trees. P1 is the taxon that diverged by reinforcement (*P. drummondii* and *P. pilosa* subsp. *pilosa*), P2 as a taxon out to P1 but not involved in the reinforcement hypothesis, P3 as the taxon sympatric with P1 driving divergence by reinforcement (*P. cuspidata* and *P. glaberrima* subsp. *interior*), and O as an individual from a proximal lineage out to all other lineages. Tests were iterated over all possible combinations of all individuals of the P1 and P3 lineages, up to three randomly sampled individuals of the P2 lineage not observed growing geographically near populations of P1 or P3, and two individuals of the O lineage. The white flowered *P. pilosa* subsp. *pilosa* lineage that evolved by reinforcement diverged from the lineage of northern *P. pilosa* subsp. *pilosa* populations (Fig. 2), therefore, we only used *P. pilosa* subsp. *pilosa* North individuals in these tests. We also considered two different species as the P2 lineage in the hypothesis concerning *P. pilosa* subsp. *pilosa* North. For all possible tests, we implemented the D-statistic test for 1,000 bootstrap iterations with loci resampling with replacement, as described in (Eaton *et al*., 2015). Tests with |Z-score| >3 are statistically significant.

We complemented the D-statistic results with TreeMix v1.13 (Pickrell & Pritchard, 2012), a model-based method for inferring population splits and migration (gene flow) under a tree framework, as implemented in iPyRAD. We targeted a potential signal of introgression by modeling a subtree of our total phylogenetic sampling. We subset the complete MS4 data matrix to include all individuals from all non-hybrid lineages in the clades containing the P1 and P3 lineages, described above, all intermediate clades, and from a lineage out to this subtree.

Individuals were grouped based on species identity with *P. drummondii* and *P. pilosa* subsp. *pilosa* North split based on flower color phenotype. We denote these subtrees as “*P. pilosa* subsp. *pilosa* North” and “*P. drummondii”* respectively. We required SNPs to be shared by at least 20% of individuals from each species and randomly sampled one unlinked SNP from each locus. The *P. drummondii* (4,638 unlinked SNPs) and *P. pilosa* subsp. *pilosa* North (3,383 unlinked SNPs) subtrees were modeled in TreeMix with *m* = 1-8 possible migration edges. The best supported number of migration edges was qualitatively chosen as the value at which the log-likelihood of the model began to increase only marginally.

### Testing origins of homoploid hybrid species

To evaluate if the five putative homoploid hybrid species in the standing *Phlox* (Table 1) have genetic support we leveraged our phylogenetic inferences and tests for gene flow to evaluate four criteria: 1) the putative hybrid sits sister to or nested within one of its putative parental taxon in our bifurcating phylogenetic reconstructions, 2) the putative hybrid shows a signal of gene flow with its other parental taxon, 3) the signal of gene flow is exclusive to the putative hybrid, and 4) the signal of gene flow goes into the putative hybrid.

For putative hybrids that are sister to or within one of their putative parent lineages, we tested for the presence of gene flow with the second putative parental lineage using Patterson’s D-statistic, as described above. For each hybrid hypothesis, a four-taxon tree was generated by subsetting the MS4 dataset to all individuals of the putative hybrid species and putative second parent species, three individuals from a species lineage between the putative hybrid and putative parent, and two individuals of a closely related outgroup. The putative hybrid and the putative parent lineages served as either P1 or P3 based on the structure of our ML phylogenies. We also considered multiple species as the P2 lineage. For putative hybrid lineages with significant evidence of gene flow with their parental taxon, we evaluated if the signature of gene flow was localized to the hybrid lineage by testing for any gene flow between the two putative parental taxa. All hybrid hypotheses tested with D-statistics were also modeled with TreeMix v.1.13, as described above. The *P. maculata* subsp. *pyramidalis*, *P. amoena* subsp. *lighthipei*, and *P. pilosa* subsp. *detonsa* modeled subtrees used 4,490, 4,294, and 1,033 unlinked SNPs respectively. We used the *P. pilosa* subsp. *pilosa* North modeled subtree for the *P. argillacea* hypothesis given their identical sampling.

### Inferring origins of allotetraploid hybrid species

We investigated the hypothesized ancestries of the four allotetraploid hybrid species (Table 1) using a consensus and comparative read alignment approach, similar to (Wang *et al*., 2021). Sequenced representatives of these taxa were previously confirmed to be tetraploid via flow cytometry at the OPGC.

First, we created reference consensus sequences from diploid lineages. Whole ddRAD loci present in at least 60% of diploid taxa from the MS4 dataset were concatenated to generate taxon-level consensus sequences using the iPyRAD window extractor toolkit, retaining the most common allele in the consensus sequence for species with multiple samples. We then mapped the filtered sequence reads for all allotetraploids to the diploid consensus sequences using bwa-mem (Li, 2013). We tallied the proportion of primary read alignments with a mapping quality ≥ 5 for the forward and reverse reads. We reasoned the diploid consensus receiving the highest proportion of quality read alignments would be most closely related to the ancestor of the mapped reads.

We tested this assumption by mapping one representative from each diploid taxon to the consensus sequences. We observed the consensus receiving the highest relative proportion of mapped reads corresponding to the known identity of the diploid individual the read came from, with some reads mapping to closely related sister taxa (Supporting Information Table S4). This method allows identification of small subclades of diploid taxa to which the reads are most similar, and we inferred the ancestry of the allopolyploid subgenomes as the diploid subclades receiving the highest rates of read mapping (Wang *et al*., 2021).

## Results

### Phylogenomic inferences of evolutionary relationships

Maximum likelihood (ML) trees inferred from the three concatenated datasets were highly congruent, despite variation in missing data (Fig. 2; Supporting Information Fig. S1, S2). Therefore, we focus our results to trends in interspecific relationships observed across phylogenetic inferences.

The base of the eastern standing *Phlox* is notably composed of long-styled taxa. We recover an early split between *P. stolonifera* and all other taxa (UFB=100), followed by a split of a well-supported clade (UFB=100) containing *P. ovata* out to *P. pulchra, P. glaberrima* subsp. *triflora*, and the allotetraploid *P. buckleyi* (UFB=100). The remaining *P. glaberrima* subspecies are found in a neighboring clade with subspecies of *P. carolina* and *P. maculata* (UFB=100). Across all inferences, the *P. glaberrima* and *P. carolina* subspecies are not monophyletic within their species delineation. The next branch subtends a small clade including *P. amplifolia* and *P. paniculata* (UFB=100).

Across all datasets, we inferred all short-styled taxa form a monophyletic group evolving from the long-styled species (UFB=100). This short-styled group contains the *P. pilosa*, *P. divaricata, P. villosissima*, and *P. amoena* subspecies, *P. floridana*, and the three Texas annual species (*P. drummondii*, *P. roemeriana*, and *P. cuspidata*), Half of our *P. pilosa* subspecies samples, including *P. pilosa* subsp. *fulgida*, *P. pilosa* subsp. *pilosa*, and *P. pilosa* subsp. *deamii*, form a clade out to the remaining short-style species (UFB=100). Given the relative northern origin of these samples, we call this the *P. pilosa* “North” group. This group also contains the white flower morph of *P. pilosa* subsp. *pilosa* that has evolved by reinforcement, sometimes described as *P. argillacea*. Both *P. divaricata* and *P. amoena* species form monophyletic groups (UFB=100), although *P. divaricata* subspecies are interdigitated.

The remaining *P. pilosa* subspecies, Texas annuals, and the allotetraploids *P. floridana, P. villosissima* subsp. *villosissma* and *P. villosissima* subsp. *latisepala* form a large clade (UFB=100). We observe reduced branch supports along the backbone of this clade; however, the Texas annual species are monophyletic (UFB=100), with *P. roemeriana* sister to *P. drummondii*. Adjacent to the annuals lie the remaining *P. pilosa* subspecies. We refer to this *P. pilosa* group as “South”, given the more southern origin of these taxa relative to the *P. pilosa* North group.

Our low support partitioning the *P. pilosa* South group from the Texas annuals may be due to the presence of the allotetraploid subspecies *P. villosissima* subsp. *latisepala* and *P. villosissima* subsp. *villosissima*. These taxa were inferred either out to the annuals or out to the *P. pilosa* South group with variable support (UFB=41-100). Variation in their inferred placement is consistent with their hypothesized hybrid ancestry between *P. pilosa* subsp. *pilosa* South and *P. drummondii* (Table 1). Similar phylogenetic uncertainty is observed for the allotetraploid *P. floridana* across analyses. This may be a result of its putative hybrid ancestry between *P. pilosa* subsp. *pilosa* South and *P. carolina* subsp. *angusta* (Table 1).

Coalescent-based species tree inference using SVDquartets recovered a species tree with nearly identical relationships to our ML inferred trees (Supporting Information Fig. S3); however, some interspecific positions varied in their branch supports. We inferred strong support for most nodes along the backbone of the tree (SBS > 80) and splits between major species groupings, except for early splits differentiating relative placement within the long-style species. *P. pulchra, P. glaberrima* subsp. *triflora*, and *P. buckleyi* remained monophyletic (SBS=84) and separate from the remaining monophyletic *P. glaberrima/P.maculata/P.carolina* subspecies (SBS=100). We also recovered the polyphyletic relationships of the *P. pilosa* North and *P. pilosa* South groups. Unlike our ML analyses, our coalescent inferences showed support for partitioning the clade of *P. pilosa* South and the allotetraploids *P. floridana, P. villosissima* subsp. *villosissima* and *P. villosissima* subsp. *latisepala* sister to the Texas annuals (SBS=89).

### Evidence of gene flow alongside reinforcement

We applied D-statistics and TreeMix v.1.13 to infer if gene flow occurred between sympatric populations of focal species in two cases of reinforcement. All D-statistic tests were structured so a negative D supports a history of introgression between these focal species.

In *P. drummondii*, reinforcement against hybridization with *P. cuspidata* has favored the divergence of light-blue to dark-red flower color in sympatry (Fig. 3a). D-statistic tests between allopatric *P. drummondii* and *P. cuspidata* show no evidence of gene flow; however, all 48 tests between sympatric dark-red *P. drummondii* and every *P. cuspidata* were negative, with 24 being significant (Table 2). Our best fit model (m=2) of the *P. drummondii* subtree in TreeMix recovered *P. cuspidata* out to *P. drummondii* and *P. roemeriana*, with a migration event going from *P. cuspidata* into dark-red sympatric *P. drummondii* (Fig. 3c). This migration edge remained present for all models m ≥2 (Supporting Information Fig. S4, S9).

**Table 2.** Summary results for four taxon D-statistic tests for introgression in two cases of reinforcement

| Reinforcement Scenario | P1 | P2 | P3 | Range D | Range Z<sup>†</sup> | nSig/N<sup>‡</sup> -D | nSig/N<sup>‡</sup> +D | Test ID |
| --- | --- | --- | --- | --- | --- | --- | --- | --- |
| *P. drummondii* | *P. drummondii* (light-blue) | *P. roemariana* | *P. cuspidata* | (-0.12, -0.01) | (0.187, 2.479) | 0/24 | 0/24 | T2:1 - T2:24 |
|  | *P. drummondii* (dark-red) | *P. roemariana* | *P. cuspidata* | (-0.16, -0.01) | (0.341, **4.692**) | **24/48** | 0/48 | T2:25 - T2:72 |
| *P. pilosa* subsp. *pilosa* North | *P. pilosa* subsp. *pilosa* North (pink) | *P. divaricata* subsp. *divaricata* | *P. glaberrima* subsp. *interior* | (-0.10, 0.08) | (0.008, 2.771) | 0/180 | 0/180 | T2:73 - T2:252 |
|  | *P. pilosa* subsp. *pilosa* North (white) | *P. divaricata* subsp. *divaricata* | *P. glaberrima* subsp. *interior* | (-0.08, 0.09) | (0.016, 2.418) | 0/90 | 0/90 | T2:253 - T2:342 |
|  | *P. pilosa* subsp. *pilosa* North (pink) | *P. paniculata* | *P. glaberrima* subsp. *interior* | (-0.09, 0.05) | (0.025, 2.707) | 0/180 | 0/180 | T2:343 - T2:522 |
|  | *P. pilosa* subsp. *pilosa* North (white) | *P. paniculata* | *P. glaberrima* subsp. *interior* | (-0.09, 0.10) | (0.026, 2.566) | 0/90 | 0/90 | T2:523 - T2:612 |

P1, P2, P3 refer to the three lineages used for the ABBA-BABA D statistic test (See Methods). Outgroups not shown. All tests can be found in Table S2.
<sup>†</sup> Z-score to assess significance of whether D deviates from zero. Bold indicates significance at $\alpha=0.01$ .
<sup>‡</sup> Number of significant tests over all possible sampled individuals. Significant -D identifies gene flow between P1 and P3, while significant +D identifies gene flow between P2 and P3.

In *P. pilosa* subsp. *pilosa* North, reinforcement against hybridization with *P. glaberrima* subsp. *interior* has favored the divergence of flower color from pink to white (Fig. 3b). Unlike reinforcement in *P. drummondii*, the ranges of *P. pilosa* subsp. *pilosa* North and *P. glaberrima* subsp. *interior* are mosaic and cannot be distinctly divided as geographically allopatric or sympatric (Levin & Kerster, 1967). Therefore, we grouped our tests for gene flow based on the flower color of the *P. pilosa* subsp. *pilosa* North individual used, pink or white, instead of by geography. Our tests show no support for gene flow between pink (N=360) or white (N=180) flowered *P. pilosa* subsp. *pilosa* North and *P. glaberrima interior* (Table 2). TreeMix modeled *P. pilosa* subsp. *pilosa* North subtrees produced similar topologies to our ML phylogenetic inferences but did not infer migration events involving *P. pilosa* subsp. *pilosa* North (Fig. 3d; Supporting Information Fig. S5, S9).

### Evidence of divergence and gene flow underlying hypotheses of putative homoploid hybrid species

We leveraged our inferred phylogenies with tests for gene flow to evaluate support for the five hypothesized cases of homoploid hybrid speciation in the eastern standing *Phlox* (Table 1). Specifically, we asked if a putative hybrid species sits nested within or sister to one of its hypothesized parental species and if a signature of gene flow exists from the other putative parental species into the putative hybrid.

Our phylogenetic inferences find support for four of the five putative hybrids being nested or within one of their putative parents (Fig. 2, Supporting Information Fig. S1, S2, S3); *P. amoena* subsp. *lighthipei* within *P. amoena* subsp. *amoena*, *P. maculata* subsp. *pyramidalis* within *P. maculata* subsp. *maculata*, *P. argillacea* within *P. pilosa* subsp. *pilosa* North, and *P. pilosa* subsp. *detonsa* in *P. pilosa* subsp. *pilosa* South. *P. pilosa* subsp. *deamii* was not found out or within *P. pilosa* subsp. *pilosa* South or *P. amoena* subsp. *amoena*, consistent with (Goulet-Scott *et al*., 2021).

For *P. amoena* subsp. *lighthipei, P. argillacea, and P. pilosa* subsp. *detonsa*, D-statistic tests and TreeMix analyses did not find support for gene flow between the putative hybrids and their second hypothesized parental species (Table 3; Supporting Information Fig. S5, S6, S7, S9). However, D-statistic tests do support gene flow between *P. glaberrima* subsp. *interior* and *P. maculata* subsp. *pyramidalis*, with 63 of 70 tests being significantly negative. Importantly, no evidence of gene flow was found between *P. glaberrima* subsp. *interior* and *P. maculta* subsp. *maculta*, N=160 (Table 3). All models of the *P. maculata* subsp. *pyramidalis* subtree in TreeMix (best fit m=4) confirmed this result, with a migration edge from *P. glaberrima* subsp. *interior* into *P. maculta* subsp. *pyramidalis* (Fig. 4, Supporting Information Fig. S8, S9). These results are consistent with *P. maculata* subsp. *pyramidalis* as a hybrid lineage arising from *P. maculata* subsp. *maculata* and *P. glaberrima* subsp. *interior* parental lineages.

**Table 3.**
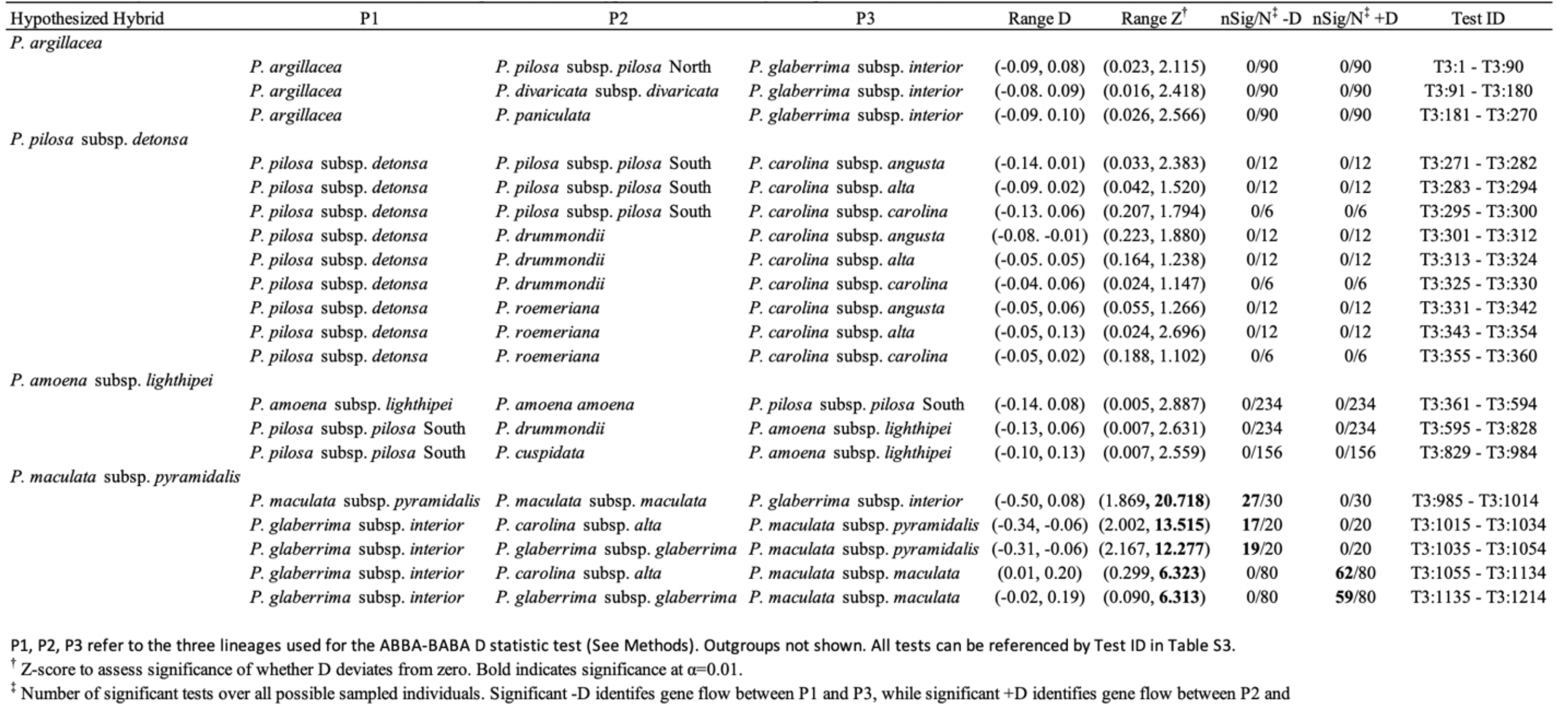
Summary results for four taxon D-statistic tests for introgression in four hypothesized cases of hybrid speciation

**Figure 4.**
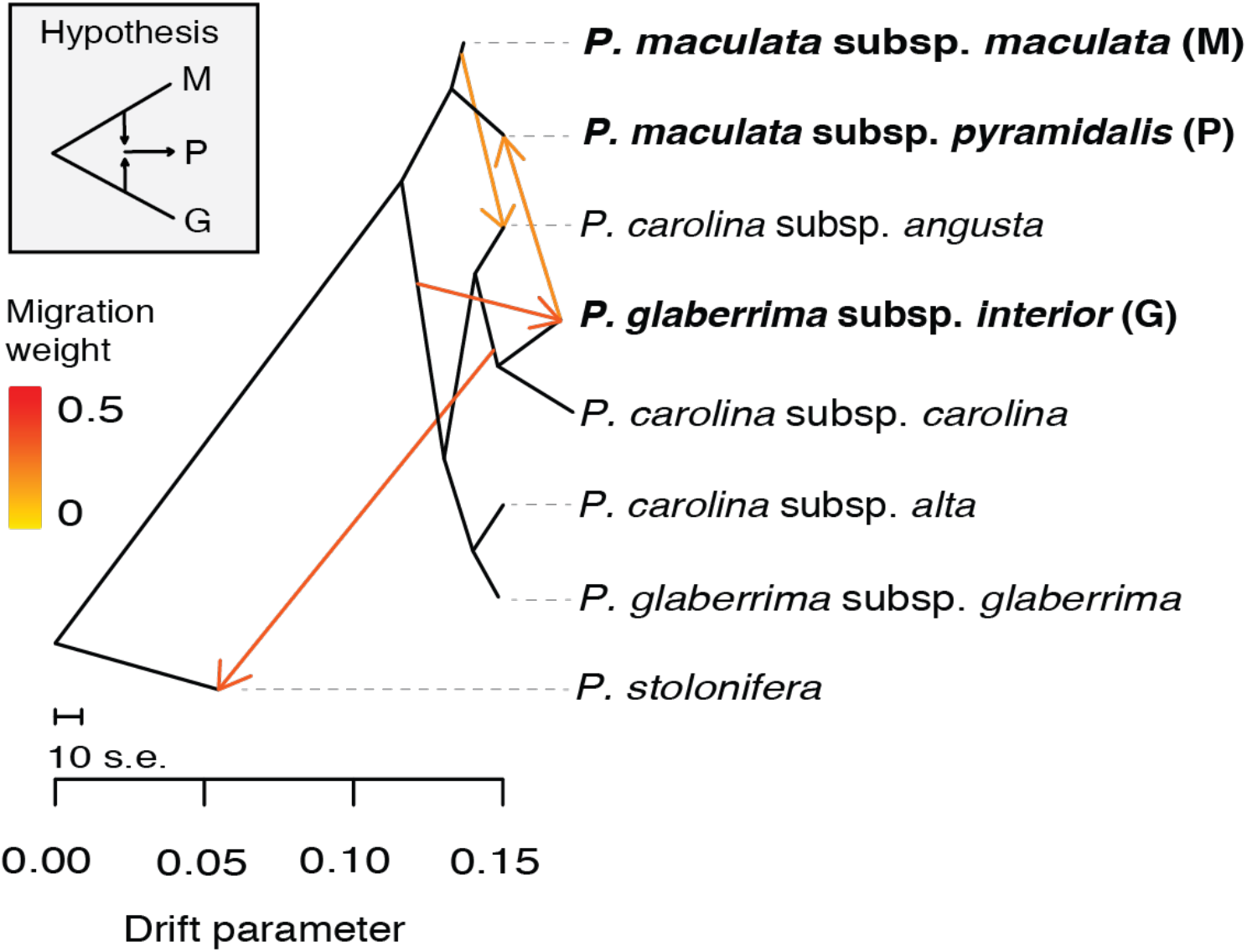
Results from TreeMix support the hybrid speciation hypothesis for *P. maculata pyramidalis* in a pruned subtree (best fit m=4). Modeling of 4,490 unlinked SNPs for individuals clustered by taxon identity inferred *P. maculata* subsp. *pyramidalis* sister to *P. maculata* subsp. *maculata* and receiving a migration event (gene flow) from *P. glaberrima* subsp. *interior*.

### Inferring origins of allotetraploid hybrid species

We leveraged comparative read mapping to infer the subgenome ancestries of four allotetraploid *Phlox* species (Table 1) (Levin, 1966b, 1968; Smith & Levin, 1967). *P. villosissima* subsp. *villosissima* and *P. villosissima* subsp. *latisepala* are thought to be independent products between *P. pilosa* subsp. *pilosa* South and *P. drummondii*. *P. floridana* is hypothesized to be the product of *P. pilosa* subsp. *pilosa* South and *P. carolina* subsp. *angusta*, and *P. buckleyi* contains two discordant diploid karyotypes hypothesized to be from extinct taxa (Wherry, 1955).

Concatenated ddRAD loci were used to generate 439,271 bp long reduced consensus sequences for each of the 28 diploid species, with a locus missingness rate of 25% among them. Mapping of reads to this consensus dataset resulted in differential assignment of sequence ancestry for each tetraploid sample to subclades of diploid taxa (Fig 5).

**Figure 5.**
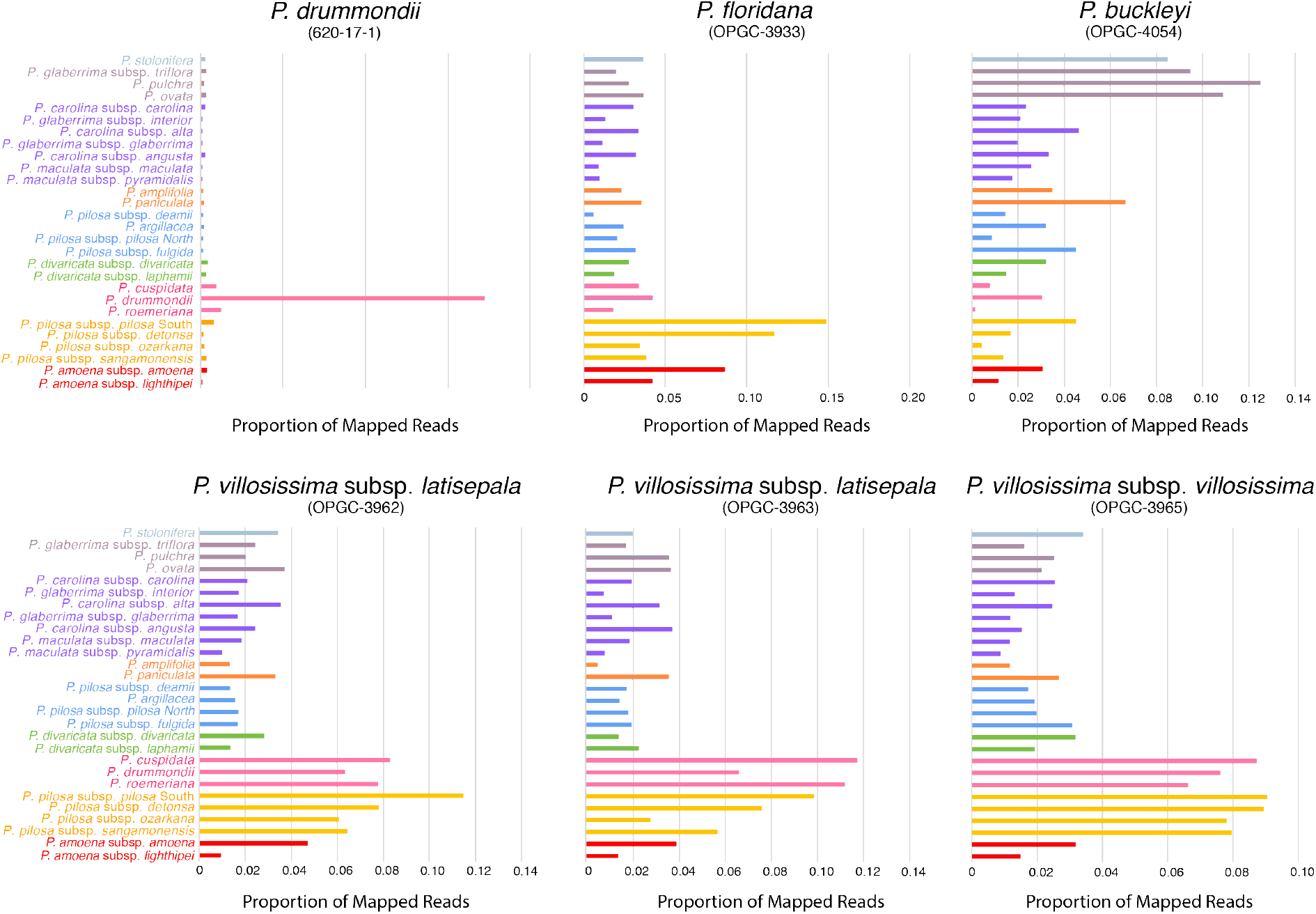
Proportion of reads mapped to diploid species-level consensus sequences for a diploid representative of *P. drummondii* and all allotetraploid hybrid individuals. Bar colors correspond to diploid species groups in Figure 2.

Reads from *P. villosissima* subsp. *villosissima* and *P. villosissima* subsp. *latisepala* mapped primarily to the *P. pilosa* South group and the Texas annuals, with each receiving 30-34% and 22-26% respectively. Within these two groups the top recipients of mapped reads were *P. pilosa* subsp. *pilosa* South and *P. cuspidata* for all three samples. *P. floridana* showed a similar high mapping rate to the *P. pilosa* South group (34%), with most reads mapping to *P. pilosa* subsp. *pilosa* South and second most to *P. pilosa* subsp. *detonsa*. Contrary to the hypothesized origins of this species, we did not observe elevated read mapping to the *P. carolina subspecies* consensus sequence. Instead, we observed elevated read mapping to *P. amoena* subsp. *amoena*. Finally, *P. buckleyi* reads mapped with the highest rate to the *P. ovata*, *P. pulchra, P. glaberrima* subsp. *triflora* clade (38%), with the highest recipients being *P. pulchra* and *P. ovata*.

## Discussion

With the accessibility of genomic data and modern phylogenomic analyses, evolutionary biology is well-positioned to understand the presence, extent, and consequences of hybridization and gene flow across large clades of organisms. Here, we present well-resolved phylogenetic relationships of the eastern standing *Phlox*, demonstrate clear support for most described species relationships, and reveal novel non-monophyletic relationships of subspecies in historically taxonomically difficult species complexes. Using this phylogenetic framework, we determine hybridization has generated new species in this clade, although maybe not as extensively as previously hypothesized. Our results support one case of hypothesized homoploid hybrid speciation, identify putative ancestries of multiple polyploid species, and find evidence of gene flow in one of two cases of reinforcement. However, we also find no support for many hypothesized cases of gene flow in the formation of new lineages. Our findings demonstrate the utility and importance of phylogenomics in confirming hypothesized evolutionary histories of non-model systems and add to the growing evidence that hybridization and gene flow across species boundaries can play diverse roles in generating novel biodiversity.

### Evolutionary relationships of the eastern standing Phlox

Previous phylogenetic inference of the eastern standing *Phlox* either relied on a handful of genetic markers or limited taxonomic sampling, leaving the evolutionary relationships of this group unclear (Ferguson *et al*., 1999; Ferguson & Jansen, 2002; Roda *et al*., 2017; Landis *et al*., 2018; Goulet-Scott *et al*., 2021). Our phylogenomic inferences on genome-wide ddRAD markers from a broad taxonomic sampling clarify the evolutionary relationships of the eastern standing *Phlox* and provide novel insight into their diversification.

Taxonomic treatments of the eastern standing *Phlox* have grouped these taxa by conspicuous differences in style length (Wherry, 1930, 1931, 1932, 1955). Unlike prior phylogenetic studies, our inferences support a large monophyletic clade of short-styled taxa subtended by multiple paraphyletic clades of long-styled taxa (Fig. 2). Divergence in style-length can generate reproductive isolation (Kay, 2006; Brothers & Delph, 2017), and this transition from long to short style-length may have reduced competition and stimulated the diversification of the short-styled clade.

Our analyses also reveal unprecedented resolution into relationships within species groups. Our inferences support *P. drummondii* and *P. roemeriana* are sister taxa within the Texas annuals (Fig. 2). This topology is supported in Roda *et al*., 2017 but not supported in studies not using genome-wide markers (Ferguson *et al*., 1999; Ferguson & Jansen, 2002; Landis *et al*., 2018). In the *P. glaberrima/P. carolina/ P. maculata* complex, we find support for non-monophyletic relationships among subspecies (Fig. 2). Additionally, the subspecies of the *P. pilosa* complex form distinct polyphyletic groups coinciding with their relative northern or southern location of origin, as suggested in Goulet-Scott *et al*., 2021. The one exception is *P. pilosa* subsp. *sangamonensis* which sits in the southern clade but is found in the north (Fig. 1), a discovery consistent with the hypothesis of *P. pilosa* subsp. *sangamonensis* arising by long-distance dispersal from southern populations of *P. pilosa* subsp. *pilosa* (Levin & Smith, 1965; Levin, 1984). The non-monophyletic relationship among the northern and southern *P. pilosa* taxa and within *P. glaberrima* were suggested in previous phylogenetic studies, but remained uncertain due to low phylogenetic supports and reliance on few genetic loci (Ferguson *et al*., 1999; Ferguson & Jansen, 2002; Landis *et al*., 2018). Only with modern phylogenomics are we able to definitively show support for these relationships.

Classic taxonomic delineations within *Phlox* are largely based on morphological characteristics (Wherry, 1955; Locklear, 2011b), yet our phylogenetic inferences demonstrate current taxonomy may not best reflect the true evolutionary relationships of some taxa within this group. Discovery of these new phylogenetic relationships begs reconsideration of the taxonomy of the group and motivates future research into why morphological variation is inconsistent with phylogenetic relationships.

### Evolution of reinforcement with and without gene flow

Limited empirical study has evaluated if interspecific hybridization generating selection for reinforcement also results in reinforcement evolving in the face of gene flow (Garner *et al*., 2018). We investigated if gene flow accompanied two cases of divergence by reinforcement in *Phlox* (Fig. 3).

*P. drummondii* has diverged from light-blue to dark-red flower color to prevent hybridization with *P. cuspidata* (Levin, 1985; Hopkins & Rausher, 2012). Our geographically widespread sampling revealed evidence of gene flow between *P. cuspidata* and multiple sympatric dark-red *P. drummondii* individuals but not between *P. cuspidata* and any allopatric light-blue *P. drummondii*, consistent with previous observations (Roda *et al*., 2017). A history of gene flow in sympatry but not allopatry mirrors the model of divergence by reinforcement to reduce hybridization in sympatry but not in allopatry (Garner *et al*., 2018). This pattern also suggests the signal of gene flow is the product of hybridization in contemporary sympatry and not due to ancient admixture between *P. cuspidata* and *P. drummondii*. Although F1 hybrids are highly sterile (Suni & Hopkins, 2018), our findings indicate hybrids do backcross in nature, resulting in interspecific gene flow. Therefore, reinforcement in *P. drummondii* evolved despite gene flow in sympatry.

Conversely, our analyses do not support a history of gene flow coinciding with reinforcement in *P. pilosa* subsp. *pilosa* North. *P. pilosa* subsp. *pilosa* North has evolved from pink to white colored flowers to prevent hybridization where it co-occurs with *P. glaberrima* subsp. *interior* in high frequency (Fig. 2) (Levin, 1966b; Levin & Kerster, 1967; Levin & Schaal, 1970a,b, 1972). Reproductive isolation between these taxa is high but field experiments and experimental crosses demonstrate these species can generate hybrid seed, with divergence in flower color reducing the formation of hybrid seed by half in nature (Levin, 1966b; Levin & Kerster, 1967; Levin & Schaal, 1970a,b, 1972)(Fig. 2). However, our analyses do not detect gene flow between these species (Fig. 3; Table 2). This result suggests hybrid offspring between these taxa may be completely inviable, lethally maladapted, or have exceedingly high sterility and that reinforcement may have occurred after gene flow had ceased.

The presence/absence of gene flow in these two cases of reinforcement present a powerful opportunity to investigate the dynamics by which reproductive trait divergence, and flower color specifically, evolves by reinforcing selection. For reproductive trait divergence to evolve by reinforcement, alleles conferring increased assortative mating within a species must remain associated with alleles causing costly hybridization between species (Servedio & Kirkpatrick, 1997; Kirkpatrick, 2000). When gene flow between species is present, recombination can disassociate these alleles, impeding successful divergence. Yet, we find the same trait evolving due to reinforcing selection in two *Phlox* species despite gene flow occurring alongside one case but not the other. This result may be the consequence of differing genetic architectures and genetic linkage underlying the cost of hybridization and flower color between these cases, varying strengths of reproductive isolating barriers between the species pairs upon their secondary contact, and/or differences in the functional cost of hybridization (i.e. offspring or gamete loss). Further study of the strength and genetic architecture of reproductive isolating barriers between these *Phlox* species will better inform how reinforcement evolves.

### Varying support for hypothesized homoploid and allotetraploid hybrid species

Within the eastern standing *Phlox* there are five hypothesized homoploid hybrid species and four hypothesized allopolyploid species (Table 1). We only observed genomic support for *P. maculata* subsp. *pyramidalis* being a homoploid hybrid species between *P. maculata* subsp. *maculata* and *P. glaberrima* subsp. *interior. P. maculata* subsp. *pyramidalis* is found phylogenetically closely related to multiple *P. maculata* subsp. *maculata* individuals (Fig. 2) with a strong signal of receiving gene flow from *P. glaberrima* subsp. *interior* (Fig. 4; Table 3.). We can conclude our sampled *P. maculata* subsp. *pyramidalis* is of hybrid origin, however future work will be necessary to discern if the hybrid origin of *P. maculata* subsp. *pyramidalis* actually gave rise to reproductive isolation of its lineage from its putative parental species (Schumer *et al*., 2018).

The remaining four hypothesized cases of homoploid hybrid speciation were not supported by our genomic investigation, despite extensive phenotypic and biomolecular evidence. In the cases of *P. amoena* subsp. *lighthipei*, *P. argillacea*, and *P. pilosa* subsp. *detonsa* the lineages thought to be hybrid in origin may simply be divergent geographically isolated populations from one of their hypothesized parental lineages. Yet in the case of *P. pilosa* subsp. *deamii*, the lineage is distantly related to both hypothesized parents and may possesses phenotypic traits resembling these species through convergence or incomplete lineage sorting.

Polyploid hybrid lineages can be significantly easier to identify than homoploid hybrids through chromosome structure and counts. Despite this advantage, inferring the evolutionary origin of these lineages remains challenging (Rothfels, 2021). Although the signals from our comparative read mapping are coarse, our approach suggests the identity of the progenitor clades that gave rise to the polyploid hybrid species in this group. Our evidence supports *P. villosissima* subsp. *latisepala* and *P. villosissima* subsp. *villosissima* are derived from *P. pilosa* (*P. pilosa* subsp. *pilosa* South) and a Texas annual species. *P. drummondii* was previously hypothesized as the parent but our analyses cannot resolve which of the Texas annual species or ancestor is the parent. As previously hypothesized, we find support for one parent of *P. floridana* to be *P. pilosa* subsp. *pilosa* South, yet we find evidence that *P. amoena* subsp. *amoena* may be the second parent instead of the *P. carolina subspecies*. We also observe *P. buckleyi* is genetically similar to the early-diverging long-styled *Phlox*, but we cannot confirm the specific parental lineages from this group. Future work using longer haplotype phase may help resolve the evolutionary origins of these species and better inform the dynamics and evolutionary trajectory of these allopolyploid species.

Across both the hypothesized homoploid and polyploid hybrid lineages, the *P. pilosa* group has been a potential and realized source of hybridization activity. Four of the five hypothesized homoploid hybrid lineages and three of the four polyploid lineages were thought to include *P. pilosa* subsp. *pilosa* as a parent. While our analyses indicate none of the hypothesized homoploid hybrid lineages have hybrid origin with *P. pilosa* subspecies, three of the four hypothesized polyploid hybrid lineages are derived from *P. pilosa* subsp*. pilosa* South *subspecies* ancestry. *P. pilosa*’s developmental biology may make it prone to evolutionary innovation, with many populations having high rates of unreduced gametes and whole genome duplications across their range (Worcester, *et al*, 2012). Future work may help address what factors underly this developmental instability and if it has an outsized role in the generation of novel evolutionary lineages relative to other *Phlox* species.

### Implications on Evolutionary History of Trait Variation

Much of the data supporting the role of hybridization and gene flow in the speciation of the eastern standing *Phlox* stemmed from comparisons of trait variation between lineages found in sympatry. Our findings support hybridization has likely contributed to some of the phenotypic variation observed across taxa in these cases, both as a source of genetic variation and force for selective divergence. However, our findings also indicate a lack of interspecific genetic exchange for many hypothesized cases of hybrid speciation. Further study of these confirmed cases may inform how specific traits and even genes move between species and contribute to reproductive isolation between lineages. While our lack of support in some cases suggests forces other than hybridization may underlie patterns of trait variation observed across taxa in this group, this finding also motivates a strong need to reevaluate our previous evidence for the existence of gene flow, and to rethink our expectations of how and why traits evolve within and between closely related species. Future investigation into the phenotypic variation of these taxa may provide new insights into how differences in ecological selective pressures across overlapping species ranges, effective population size, and incomplete lineage sorting contribute to phenotypic divergence within and between species.

## Acknowledgements

We are grateful to S.V. Edwards, E.M. Kramer, J. Wakeley, and J. Mallet for thoughtful discussions that improved this study, and to C.J. Ferguson, G.A. Burgin, and A.E. Berardi for their helpful comments on this manuscript. We are also thankful to P.S. Jourdan and the OPGC staff for sharing their *Phlox* collections with us.

## Author Contributions

A.G.G. and R.H. conceptualized the project. A.G.G., B.E.G.S., and R.H. collected and analyzed the data. A.G.G. and R.H. wrote the manuscript.

## Data Accessibility

Supporting data underlying this article is found on Dryad Digital Repository: [insert rep ID}

## Supplementary Information

**Figure S1.** Maximum likelihood phylogenetic inference from whole concatenated loci, with a locus shared among at least 10 individuals.

**Figure S2.** Maximum likelihood phylogenetic inference from whole concatenated loci, with a locus shared among at least 20 individuals.

**Figure S3.** Coalescent-based phylogenetic inference using SVDquartets, with a locus shared among at least 30 individuals.

**Figure S4.** TreeMix models for the *P. drummondii* subtree.

**Figure S5.** TreeMix models for the *P. pilosa subsp. pilosa* North subtree.

**Figure S6.** TreeMix models for the *P. pilosa subsp. detonsa* subtree.

**Figure S7.** TreeMix models for the *P. amoena subsp. lighthipei* subtree.

**Figure S8.** TreeMix models for the *P. maculate subsp. pyramidalis* subtree.

**Figure S9.** TreeMix model best fit estimation.

**Table S1.** Species sampling.

**Table S2.** All D-statistic test results for introgression in two cases of reinforcement.

**Table S3.** All D-statistic test results for introgression in four hypothesized cases of hybrid speciation.

**Table S4.** Summary of comparative read mapping of diploid and allotetraploid species to diploid consensus sequences.

