## Supporting Information for "Phylogenomic patterns of divergence and gene flow detail the evolution of reinforcement and hybrid speciation in *Phlox* wildflowers"

**Fig. S1:** Maximum likelihood phylogenetic inference of eastern standing *Phlox* relationships from whole concatenated ddRAD loci. Loci were required to be shared by at least 10 individuals. Numbers on branches represent bootstrap supports from 100 bootstrap replicates using ultrafast bootstrap approximation (UFB). Bootstrap supports are shown for all nodes differentiating between taxa.

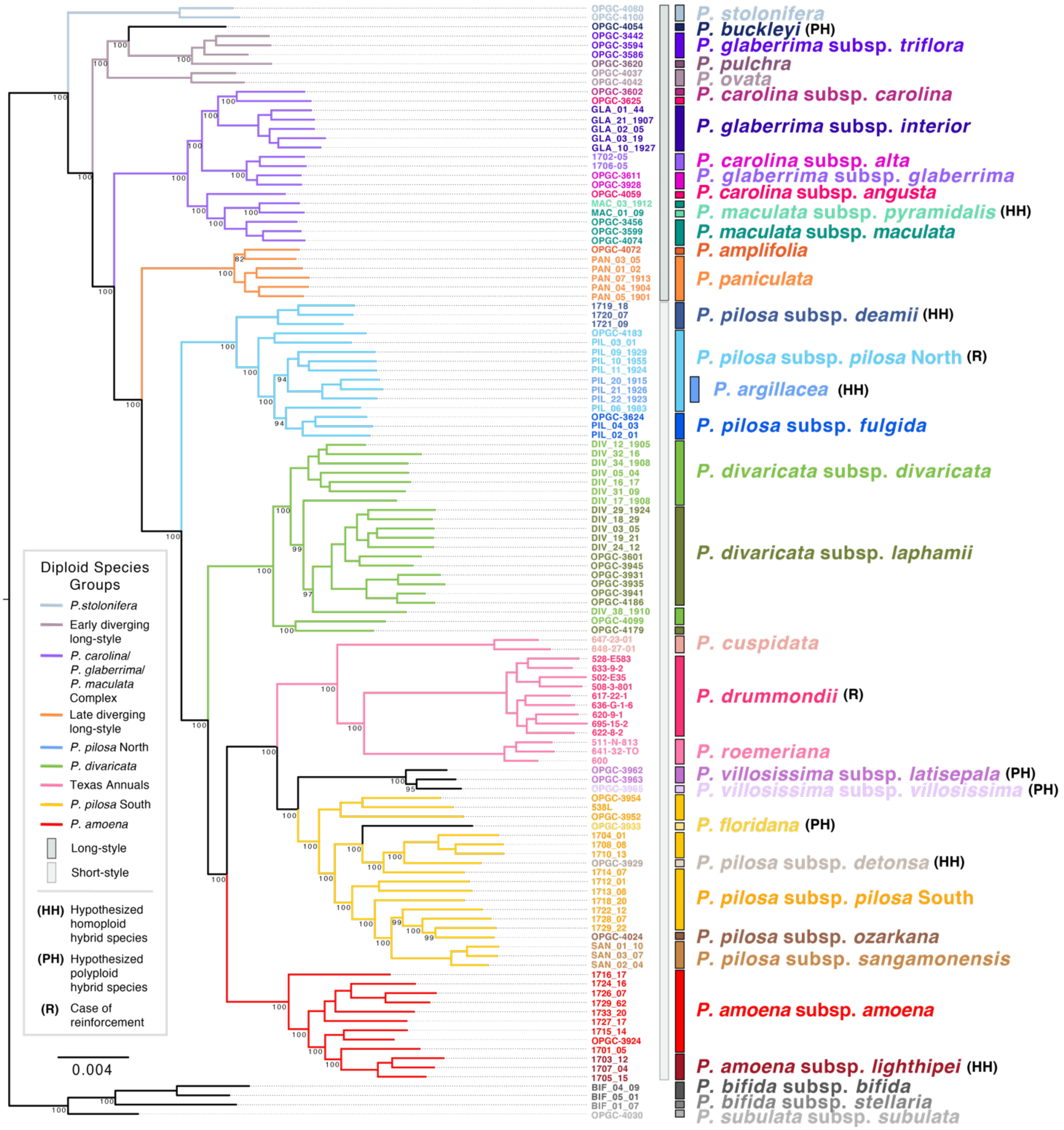

**Fig. S2:** Maximum likelihood phylogenetic inference of eastern standing *Phlox* relationships from whole concatenated ddRAD loci. Loci were required to be shared by at least 20 individuals. Numbers on branches represent bootstrap supports from 100 bootstrap replicates using ultrafast bootstrap approximation (UFB). Bootstrap supports are shown for all nodes differentiating between taxa.

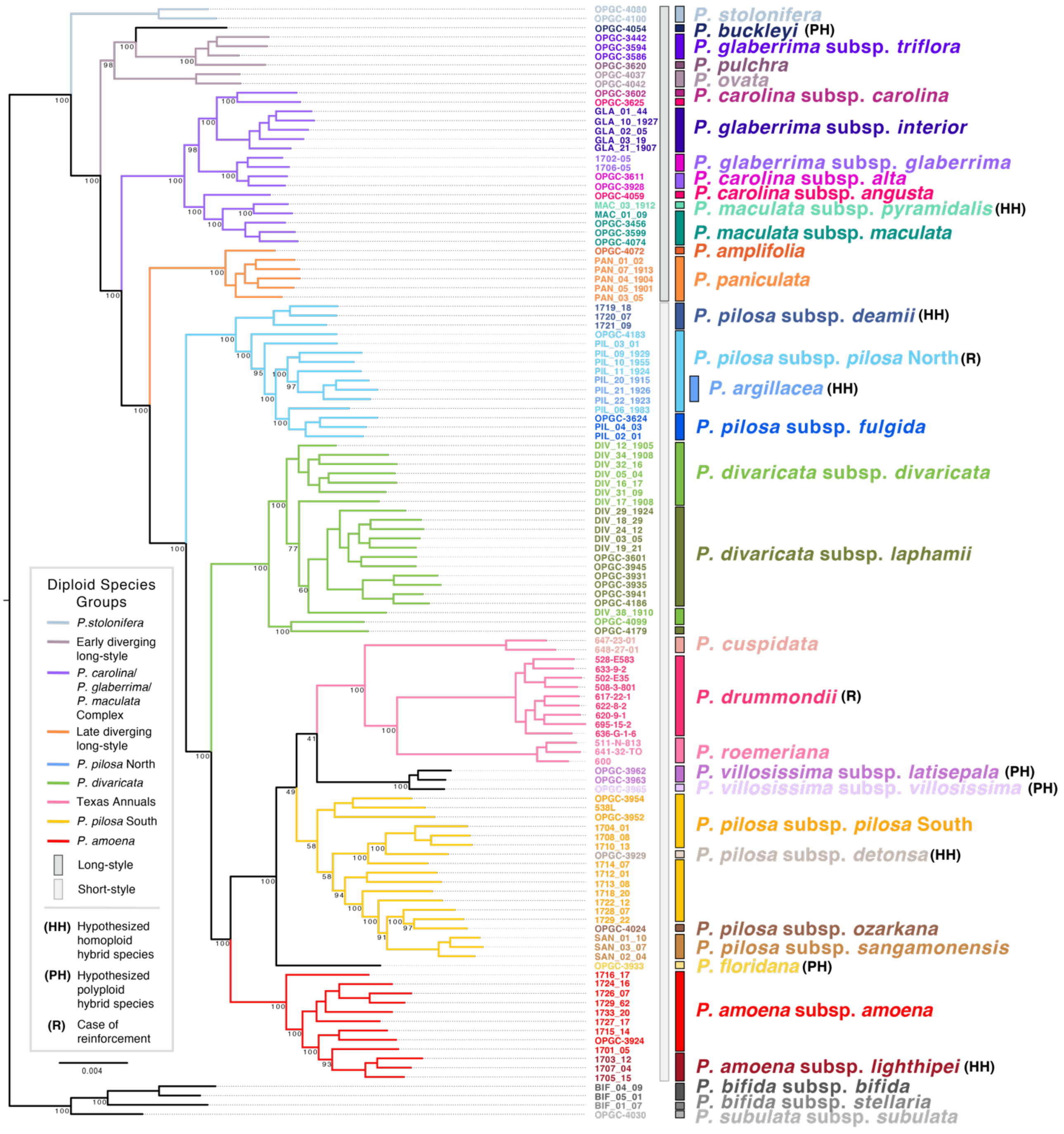

**Fig. S3:** Coalescent-based phylogenetic inference using SVDquartets of eastern standing *Phlox* relationships from 5,306 unlinked SNPs. Loci were required to be shared by at least 30 individuals. Numbers on branches represent bootstrap supports from 100 nonparametric bootstrap replicates (SBS). Unlabeled branches had supports of SBS=100.

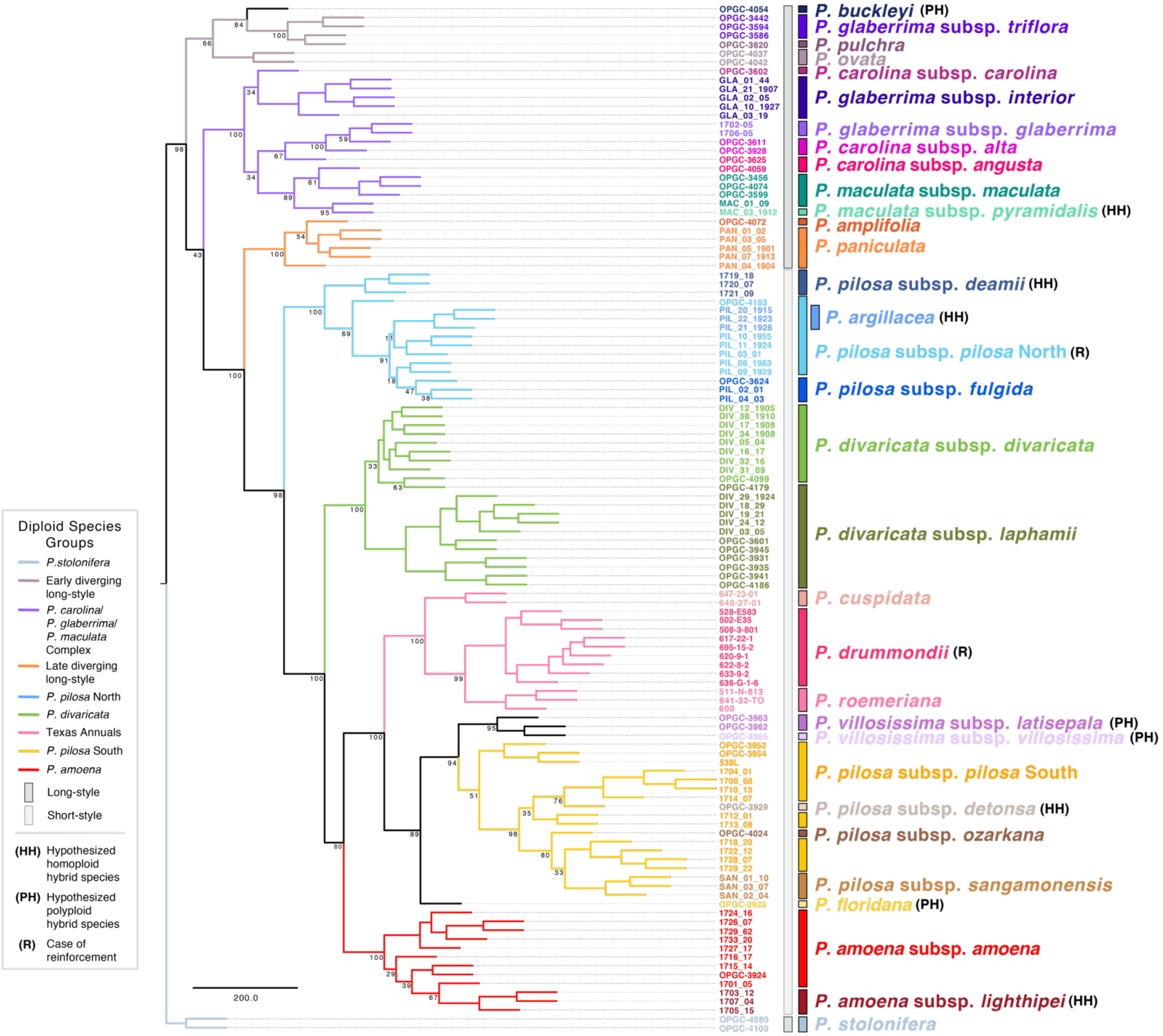

**Fig. S4:** TreeMix models for the *P. drummondii* subtree with m=1-8 migration events. Best fit model (m=2) identified by the decay in log likelihood across models (Supporting Information Fig. S9).

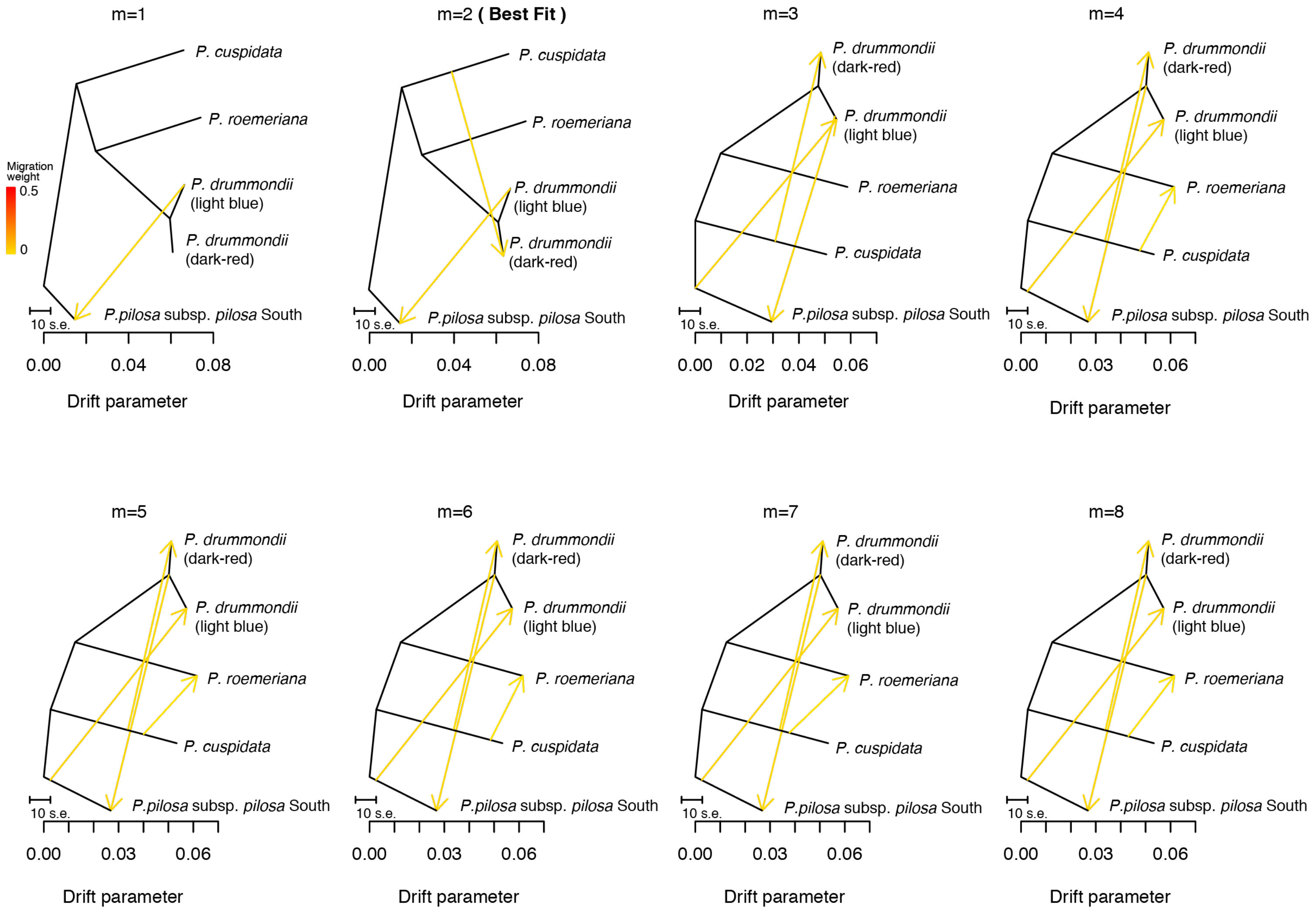

**Fig. S5:** TreeMix models for the *P. pilosa* subsp. *pilosa* subtree with m=1-8 migration events. Best fit model (m=1) identified by the decay in log likelihood across models (Supporting Information Fig. S9).

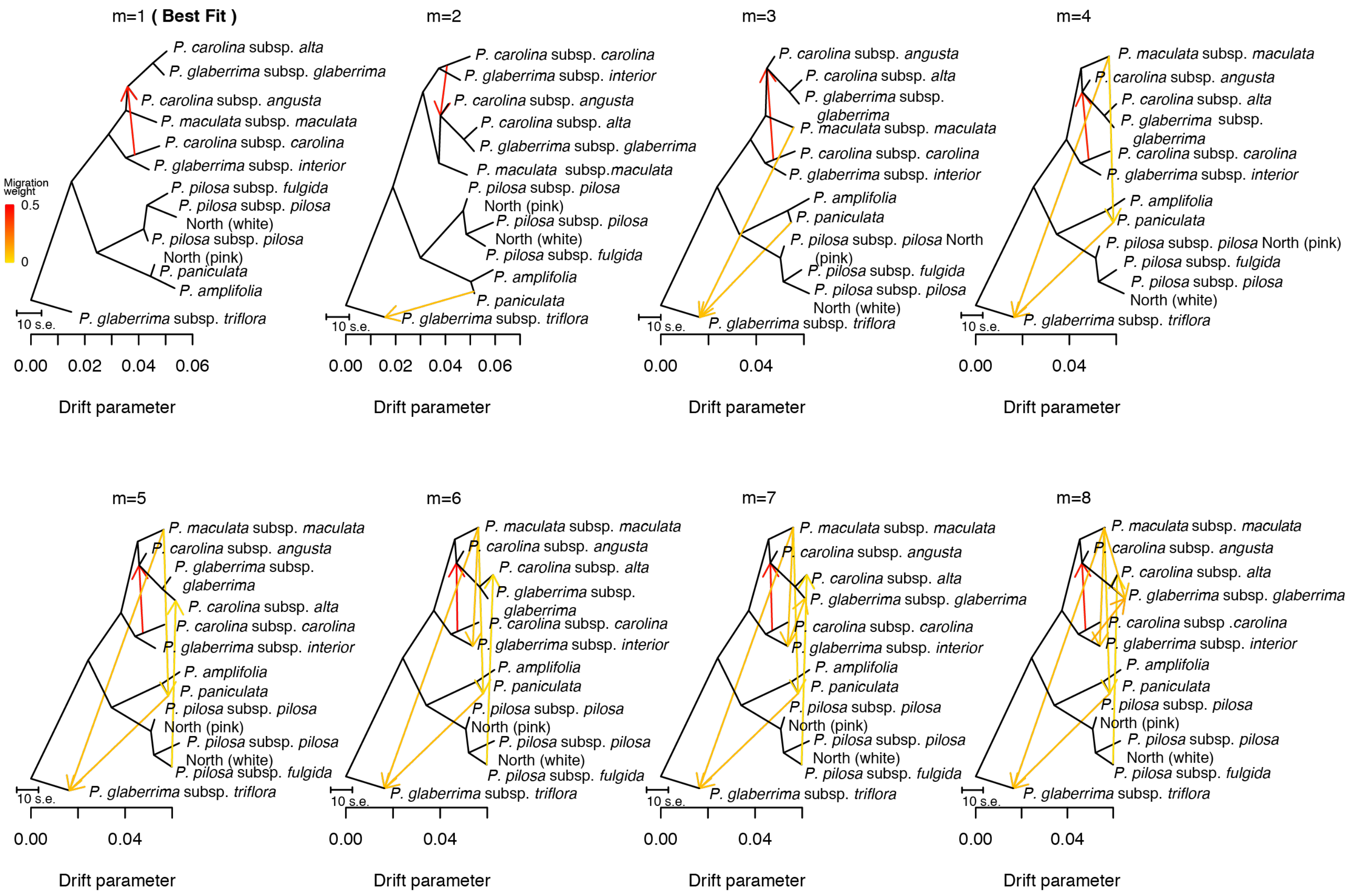

**Fig. S6:** TreeMix models for the *P. pilosa* subsp. *detonsa* subtree with m=1-8 migration events. Best fit model (m=3) identified by the decay in log likelihood across models (Supporting Information Fig. S9).

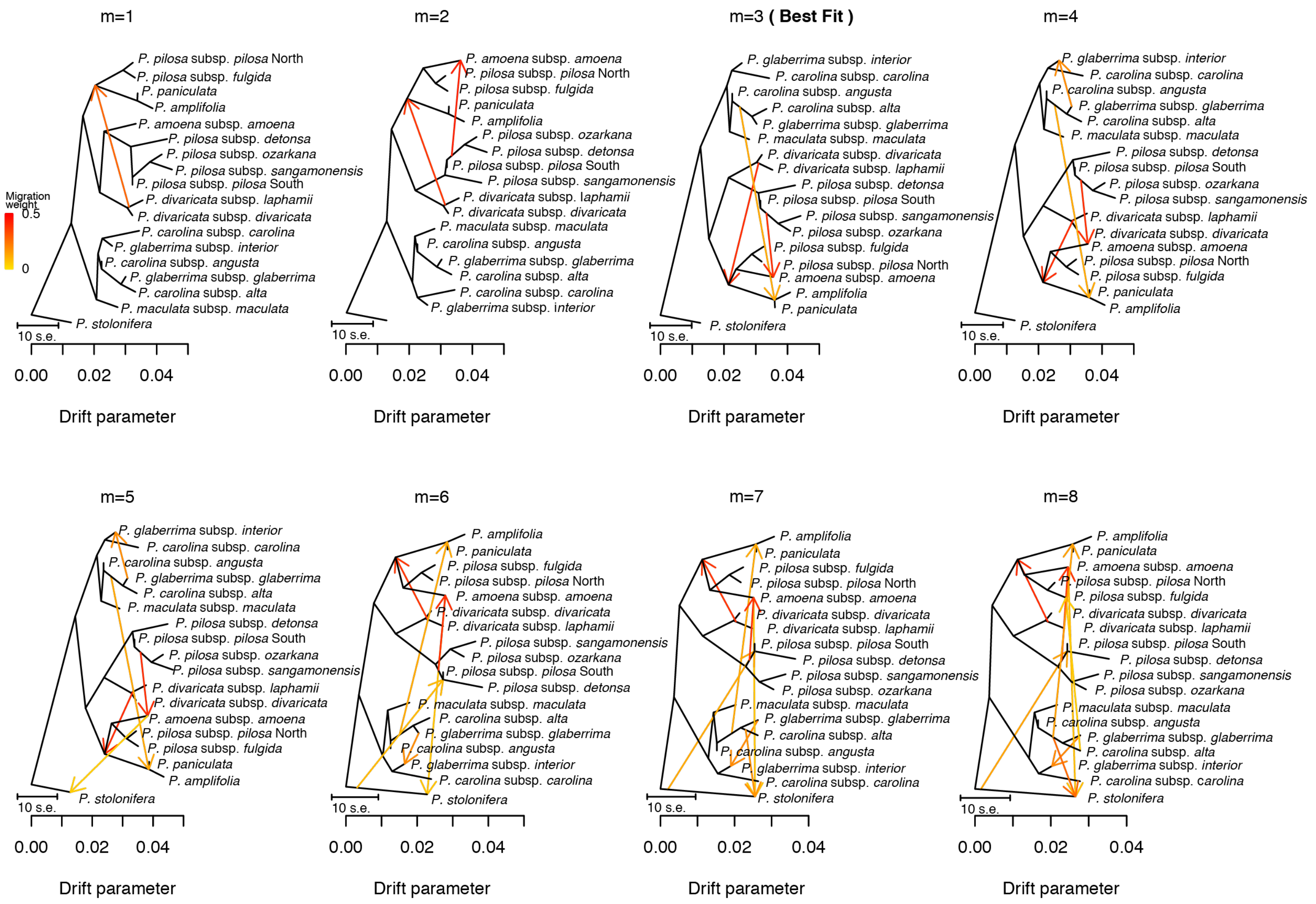

**Fig. S7:** TreeMix models for the *P. amoena* subsp. *lighthipei* subtree with m=1-8 migration events. Best fit model (m=1) identified by the decay in log likelihood across models (Supporting Information Fig. S9).

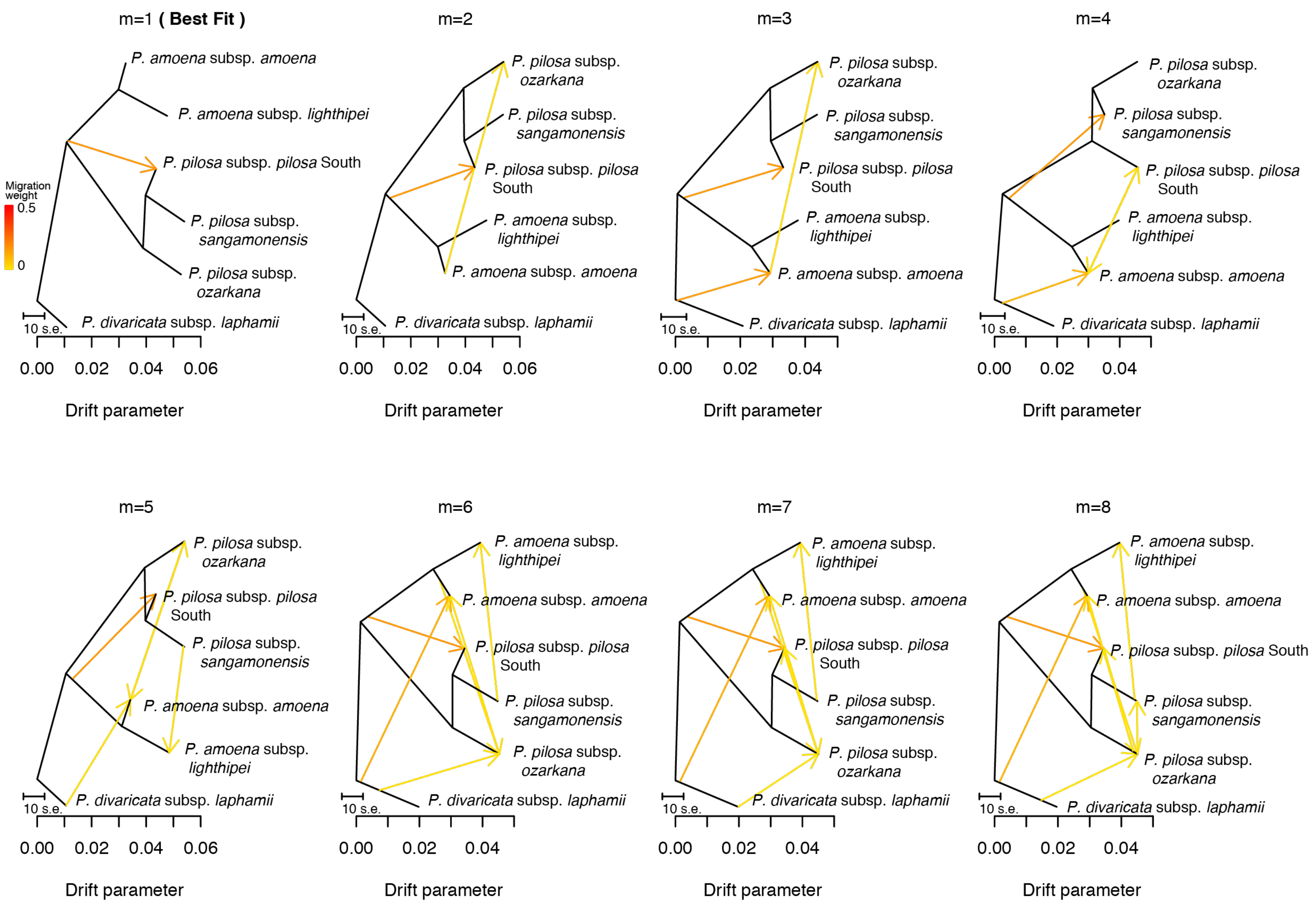

**Fig. S8:** TreeMix models for the *P. maculata* subsp. *pyramidalis* subtree with m=1-8 migration events. Best fit model (m=1) identified by the decay in log likelihood across models (Supporting Information Fig. S9).

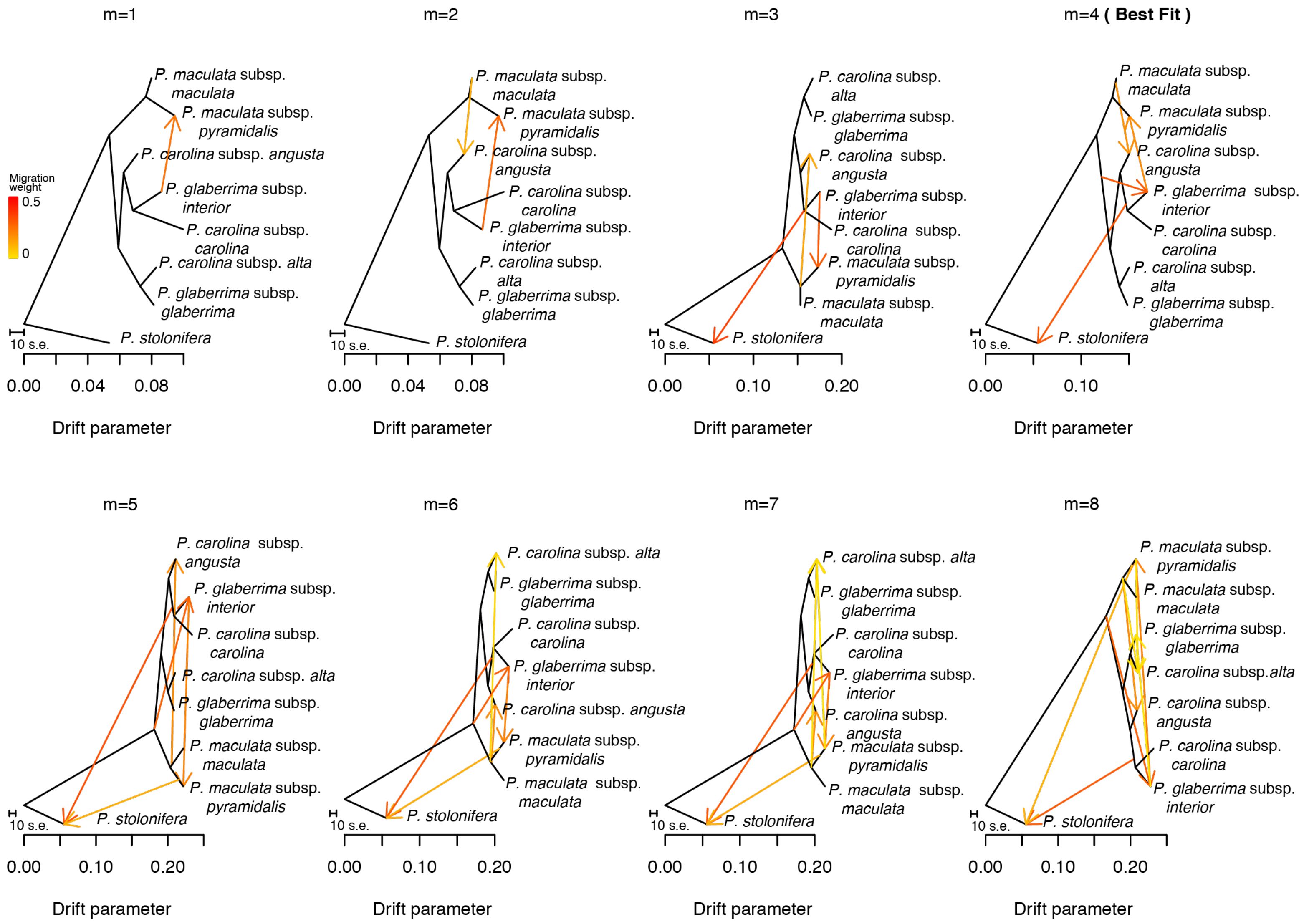

**Fig. S9:** Qualitative approximation of best fit TreeMix model by the decay in increase in log likelihood across trees modeled with TreeMix with increasing number of migration events m = 0-8.

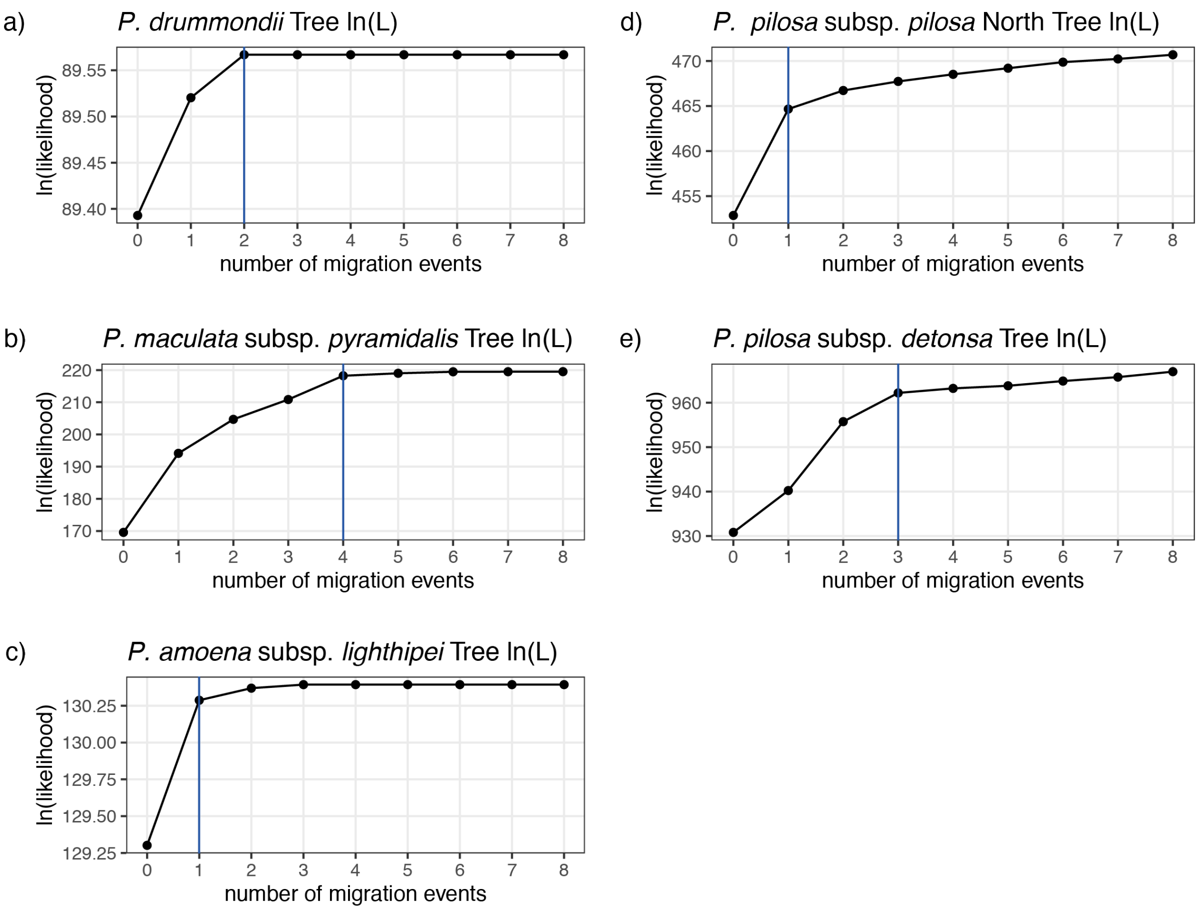
